# Establishing a co-culture aggregate of N-cycle bacteria to elucidate flocculation in biological wastewater treatment

**DOI:** 10.1101/2024.07.02.601720

**Authors:** Laurens Parret, Kenneth Simoens, Benjamin Horemans, Jo De Vrieze, Ilse Smets

## Abstract

Biological flocculation is a complex phenomenon that is often treated as a black box. As a result, flocculation problems are usually remediated without knowledge of the exact causes. We show that it is feasible to exploit a model (N-cycle) consortium with reduced complexity to fundamentally study bioflocculation. Strong nitrifier microcolonies were formed during oxic/anoxic cycles in sequencing batch reactors, using alginate entrapment as a cell retention system. After release of these aggregates into suspension, macroclusters with flocs of the denitrifier were observed. These results suggest that a living model of a full-scale activated sludge floc can be built through the use of this bottom-up approach. By eliminating shifts in the microbial community, the applied experimental conditions have a more direct effect on the observations.

## 1. Introduction

In engineered systems for biological wastewater treatment (and resource recovery), the importance of natural aggregation is widely accepted (Srivastava et al. (2018)). Biological flocculation not only yields a clear effluent, it can intensify (granular) sludge separation, and retains functional microorganisms in the system (Chen et al. (2020)). However, the sludge itself is mostly still treated as a black box (Suresh et al. (2018)). Especially the matrix of extracellular polymeric substances (EPS) in which the microorganisms are embedded, is a complex structure that should be more elaborately investigated to be able to link it to functionality (Felz et al. (2019)). Unravelling EPS is a difficult task, given its variability between biofilms (and microorganisms) and the challenges (inaccurate colorimetric analysis and interferences) that arise when analysing the biopolymers, (Flemming and Wingender (2010); Lin et al. (2018)). As a result of this lack of insight, EPS-related flocculation problems are usually still remediated ad-hoc in practice (Wang et al. (2018)), and mechanistically controlled flocculation (or granulation) is not yet possible.

To help unravel the full-scale complexity of activated sludge, large scale 16S rRNA gene sequencing approaches, such as the MiDAS project (Dueholm et al. (2023)) have provided insight into the communities that shape global wastewater treatment systems. The result hints towards a common trend in abundant species (Dueholm et al. (2022)). Functionally relevant organisms have been extensively visualised by fluorescent in-situ hybridisation (FISH) to obtain information about their spacial vicinity (Per Halkjær Nielsen and Lemmer (2009)). Key species of for example ammonium- and nitrite oxidising bacteria (AOB and NOB) have been isolated and studied individually for decades (Laanbroek et al. (2002); Nogueira and Melo (2006)). Knowledge of these underlying biological processes may contribute to solving more large-scale problems, as shown by Mellbye et al. (2018) for nitrogen oxide emissions.

To our knowledge, there is no study that exploits the monitoring potential of pure cultures in a model consortium of activated sludge to fundamentally study bioflocculation. Working with more defined communities would drastically reduce the complexity of advanced monitoring tools, with “omics” techniques as the final destination (for example already done for AOB by Yu et al. (2018)). Building model consortia has already been extensively described in biofilm research to simulate important properties of the real system, from the medical field (Ghesquière et al. (2023)) to pollutant degradation in soil (Horemans et al. (2014)), all indicating the power of mixed-species biofilms (Strathmann et al. (2000)). As summarised by Hobley et al. (2015), the current advancements in monitoring tools provide more fundamental knowledge on the formation of (both beneficial and pathogenic) mixed-species communities. In the wastewater treatment context, multi-species interactions in the nitrogen cycle are mostly examined in terms of metabolite consumption and spacial distribution (Pérez et al. (2014); Montràs et al. (2008)), or combined with modelling (Cruvellier et al. (2016)).

This study aims to provide a proof-of-concept toolbox to form flocs (spherical biofilms) *de novo*, starting from selected N-cycle microorganisms. By growing them together, the microenvironment to which they are exposed in a real system is mimicked. More controlled experimental boundary conditions can then be applied to obtain a better understanding of the fundamental requirements for bioflocculation. After all, top-down approaches on sludge, such as bioaugmentation, are challenging and often lead to inconclusive results (Herrero and Stuckey (2015); Christiaens et al. (2023)). Hence, the hypothesis of the proposed bottom-up approach was that the experimental results can be more directly related to the applied operational conditions by eliminating shifts in the microbial community.

To improve retention of the slow-growing nitrifiers, cell immobilisation through (reversible) alginate bead entrapment is employed during startup. Hydrogel entrapment is a proven method in many fields, with encapsulated cells ranging from complete activated sludge (Bouchez et al. (2009)) or enriched flocs (Hill and Khan (2008)), to stem cells (Tang et al. (2012)). Recently, in the work of Samarasinghe et al. (2016), enhanced nitrification was even observed in their “environment cubes” by immobilisation of AOB and NOB. A more complete overview of the potential of encapsulation is given for example by Wang et al. (2021). Interestingly though, the recent work of Pabst et al. (2016) (in the medical field), suggests that the artificial encapsulation of *Staphylococcus aureus* provided the necessary matrix to form bacterial aggregates that mimic the real infection system, despite the exogenous nature of the hydrogel. In essence, that is exactly what we aim for as well, though in a wastewater treatment context. Hence, the second hypothesis was that the reduced community can still perform the same functionality in a standard oxic/anoxic cycling environment, and flocculate accordingly given the appropriate conditions (e.g., feast-famine (Sun et al. (2021))). Through encapsulation, the microorganisms are not only retained in the system, but also forced together in the polymer matrix.

## 2. Materials and methods

### 2.1. Experimental overview

In this work, both entrapped and planktonic experiments were performed with different biomass content. The main research structure is summarised in Figure 1.

**Figure 1:**
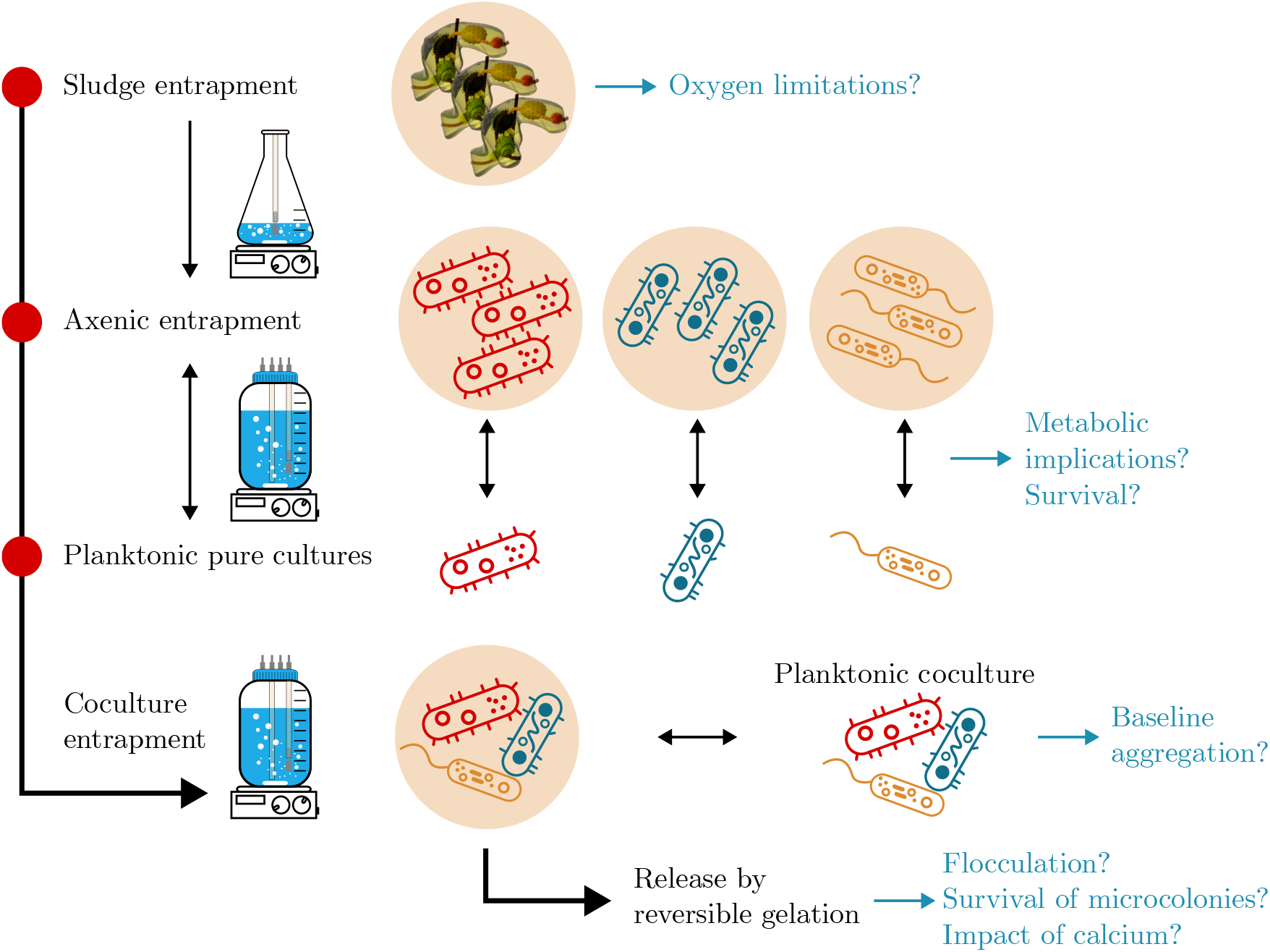
Schematic overview showing the interactions and main research goals of the performed experiments. Both entrapped and planktonic reference experiments support the main coculture entrapment (and release) experiment.

### 2.2. Microbes and media

The applied N-cycle bacteria are illustrated in Figure 2. The ammonium oxidising bacterium (AOB) *Nitrosomonas europaea* DSM 28437 and nitrite oxidising bacterium (NOB) *Nitrobacter winogradskyi* DSM 10237 were obtained from DSMZ (Leibniz Institute DSMZ - German Collection of Microorganisms and Cell Cultures GmbH). A mutant strain of the denitrifying bacterium (DEN) *Azoarcus communis* Swub 3 DSM 12120, expressing the fluorescent protein mScarlet-I (*A. communis* Rif^*r*^ -mSc) was used, as described by Christiaens et al. (2023). The preliminary entrapment test with activated sludge (to test oxygen limitations) was performed in stirred erlenmeyer flasks with the same medium and biomass as in Christiaens et al. (2022). Precultures of AOB and NOB were shaken at 28 °C (in the dark) in a phosphate-buffered coculture medium with ammonium and nitrite respectively (summarised in supplementary Table S1). Carbonate was added and yeast extract was left out to promote autotrophic growth. For AOB, the pH was corrected during growth using 5 wt% NaHCO_3_ when the medium turned green (cf. DSM medium 1583). *A. communis* Rif^*r*^ -mSc was first grown on Tryptic Soy Agar (TSA) with 20 mg L^−1^ rifampicin and 20 mg L^−1^ gentamicin and precultured in Tryptic Soy Broth (TSB, Oxoid). For experiments with entrapped biomass, the reactors were operated at 28 °C with HEPES-buffered medium. To dissolve the alginate, the supernatant was replaced with P-buffered medium to extract and precipitate all calcium, which was acting as crosslinking agent in the hydrogel. The detailed composition of the medium can again be found in supplementary Table S1.

**Figure 2:**
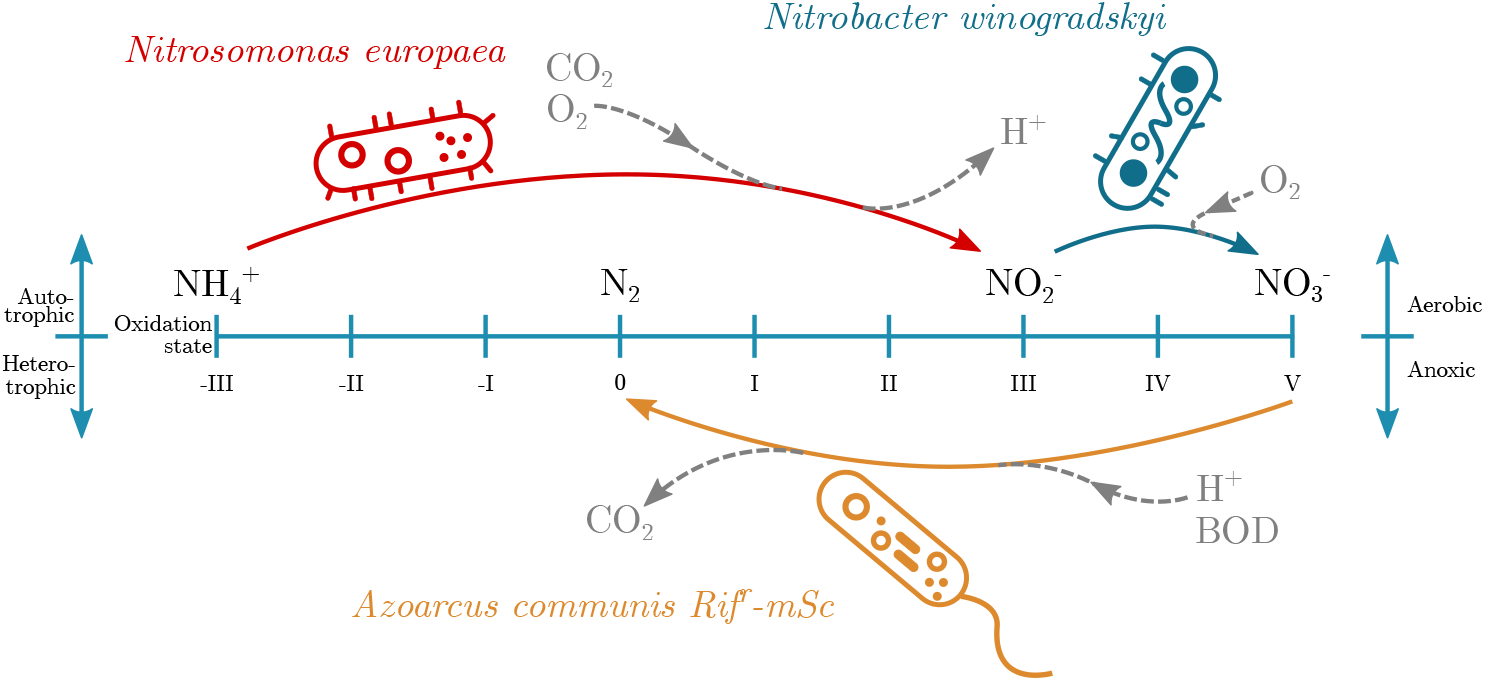
Applied bacteria and their position in the nitrogen cycle. Figure inspired by Canfield et al. (2010).

Cell counts were determined by flow cytometry using the Cytek Amnis CellStream and SYTO 9 (Invitrogen) at a final concentration of 1.67 µM in Phosphate Buffered Saline (PBS). Gating was performed on the emission channel corresponding to the 488 nm laser, and expression of mScarlet-I was confirmed for *Azoarcus* in the 561 nm channel. The effect of noise (e.g. from salts) could be reduced by rejecting all events with an aspect ratio in the side scatter (SSC) channel equal to 0 or 1.

### 2.3. Sterile entrapment and reactors

Sodium alginate powder (Alpha Aesar, B25266 low viscosity) was sterilised by thoroughly spreading it out on autoclaved stainless steel instrument trays (Rotilabo, 36 cm × 24 cm), and applying UV irradiation from a Biovanguard 4 (Telstar) UV-C lamp (30 W), twice for 30 min with intermittent manual shaking of the dusted tray. A maximum of 1 g was loaded on each tray at a time (±1.5 mg/cm^2^) to improve the uniformity of the powder layer. Sterile alginate was harvested using cell scrapers, dissolved in distilled water overnight and diluted to a concentration of 1.5 wt% after biomass addition. All bacteria were added to sodium alginate with a final concentration of 5 × 10^6^ cells/mL.

Polymer beads were produced using the Nisco VARD2Go jet-breakup encapsulation unit at 1.5 barg and 0.42 kHz with a 500 µm sapphire nozzle. Gelation occurred in an excess of autoclaved 5 wt% CaCl_2_ during stirring for at least 10 min. The encapsulator was placed inside a plexiglass box cleaned with Umonium (Huckert’s). To ensure a positive outward pressure, two outlets of the Nisco pressure control unit (setpoint 0.8 barg) were connected to Sartorius Midisart 2000 PTFE Air Filters on the box, leading to a laminar-like flow inside the cabinet. A schematic overview of the setup is shown in Figure 3a, along with a picture in supplementary Figure S1.

**Figure 3:**
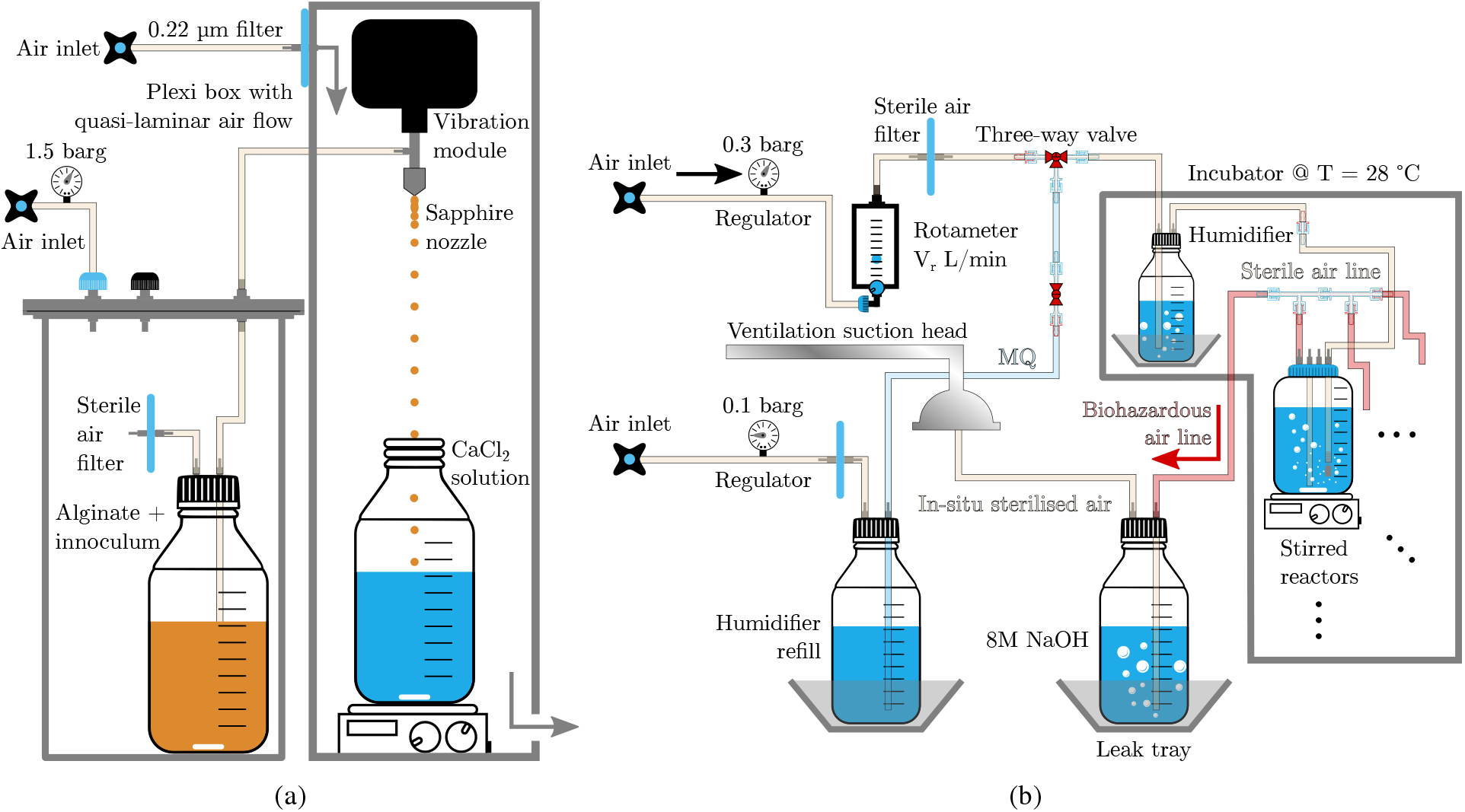
Schematic overview of the a) axenic bead production setup b) applied bioreactor design.

Beads with entrapped biomass were washed three times with autoclaved demineralised (demi) water to remove the CaCl_2_ and resuspended in coculture medium at a ratio 1:10 by volume. Hence, the final cell number for each strain was 5 × 10^5^ cells/mL in the reactors. Medium in (planktonic) reference reactors was diluted with demi water to obtain the same initial ion concentrations. Reactor vessels were operated as illustrated in Figure 3b (see supplementary Figure S2 for photos). The humidification system (with Milli-Q (MQ) water) serves to prevent volume losses due to the sparging of dry compressed air or nitrogen gas. Stainless steel sintered aeration stones (Brouwland, for wort aeration) were applied to generate small gas bubbles in the reactor. During oxic conditions, a flow rate of 1 vvm (gas volume per broth volume per minute) was applied, while anoxic conditions were generated by sparging N_2_ at 1 vvm for 5 min, and at 0.25 vvm for the remaining part of the phase (to prevent clogging of the aerator and maintain reactor overpressure). A phase switch from oxic to anoxic conditions (or vice-versa) was performed either when all ammonium (or nitrate) and nitrite was depleted, or when the values stayed constant for 1-3 days (qualitatively, using Quantofix (Macherey-Nagel) strips). Nitrogen and carbon sources were spiked in the coculture medium (to not dilute the trace elements and to compensate for volume losses due to sampling) and added (at these switch times) through a Filtropur 0.22 µm filter attached to the reactor lid.

### 2.4. Microscopy, cryosectioning and qPCR

Samples of alginate beads were taken by withdrawing 10 mL from the reactor to obtain approximately 1 mL of beads. The supernatant was sterile filtered and injected back into the reactor. Half of the sampled beads were fixed in 4 % PFA (see Per Halkjær Nielsen and Lemmer (2009)) at 4 °C overnight, washed three times with demi water (instead of Phosphate Buffered Saline (PBS) to preserve bead integrity), and incubated overnight at 4 °C in OCT (Optimal Cutting Temperature) compound (VWR) before storage at −20 °C for cryosectioning and fluorescent in-situ hybridisation (FISH). The leftover beads were weighed on an analytical balance, and washed three times with demi water. To dissolve them and preserve the DNA content, a citric acid-EDTA buffer was added as described by Lopez et al. (2017). The suspension was stored at −20 °C for qPCR analysis.

Bright field (BF), phase contrast and fluorescent images were captured using the Olympus IX83 inverted microscope. To enhance BF contrast, complete beads were stained with crystal violet and washed with demi water before acquisition.

Confocal images were obtained using an Olympus Fluoview FV 1000 CLSM. Fluorescent in-situ hybridisation (FISH) on suspended samples was performed as described by Per Halkjær Nielsen and Lemmer (2009). An overview of the applied FISH probes is provided in supplementary Table S2. The PFA fixation and dehydration steps of the protocol from this handbook were also applied as pretreatment steps for Scanning Electron Microscopy (SEM) using the JSM-6010LV (JEOL), with the exception that absolute ethanol (instead of 96%) was applied in the final dehydration step before drying at 46 °C. Samples were then coated with a Au/Pd layer (JEOL JFC-1300 Auto Fine Coater, for 30 s at 30 mA under Ar plasma) before imaging.

Cryosections of alginate beads were obtained by first embedding the sample in between layers of frozen OCT in 35 mm petri dishes. The resulting disk was transferred to a specimen chuck of the NX70 cryostat and 50 µm slices were cut at a blade and specimen temperature of −10 °C and −15 °C, respectively. Sections were immobilised on Superfrost Plus (Menzel Gläser) slides and stored at −20 °C. For FISH on cryosectioned samples, the slides (containing sections and therefore fixed biomass) were coated with agarose by dipping the slide into a 50 mL falcon with molten 1 wt% agarose (at ±40 °C) for a few seconds, and freezing it in place by cooling the slide on a freezer element at −20 °C. This procedure is described in Chapter 8 of the FISH handbook (Per Halkjær Nielsen and Lemmer (2009)) to improve retention of sludge samples. In case of the hydrogel sections, the agarose preserves their integrity and avoids complete loss of the sample when the OCT compound is removed during dehydration (dissolves in ethanol), and when the hydrogel weakens in the washing buffer (contains only monovalent ions).

The DNA extraction was performed using the QIAamp DNA Mini Kit (Qiagen) following the instructions of the manufacturer. Oligonucleotide design, acquisition of template DNA for qPCR standards, and initial SYBR Green (Thermo Scientific ABsolute QPCR Mix) assays for validation were performed as described by Horemans et al. (2016). More information on the standard templates as well as (q)PCR conditions and oligonucleotides can be found in supplementary information Section S.1.2. All samples were run in multiplex (FAM, JOE, ROX) TaqMan assays on a Rotor Gene Q (Qiagen) using Takyon 2X MasterMix (Eurogentec) in triplicate. Tenfold dilutions of the extracted DNA samples were also analysed to verify the absence of inhibitory effects in the PCR reaction.

### 2.5. Metabolite measurement

All samples for extracellular metabolites were sterile filtered using Filtropur 0.22 µm syringe filters. Anions and cations were determined by ion chromatography (IC) using conductivity detection. The anion IC (Metrohm Eco IC) was equipped with a chemical suppressor (regenerated with 500 mM H_2_SO_4_ and 500 mM oxalic acid), a Metrosep A Supp 17-250/4.0 column, and eluent containing 0.2 mM NaHCO_3_ and 5 mM Na_2_CO_3_. Another Metrohm Eco IC was connected in series for the simultaneous analysis of cations in the same sample, using a Metrosep C6-100/4.0 column and eluent containing 1.7 mM HNO_3_ and 1.5 mM dipicolinic acid. Both systems had a 10 µL sample loop and a quantification limit of 0.5 ppm. Glucose, acetate and lactic acid concentrations were measured on an Agilent 1200 HPLC, fitted with a Aminex HPX-87 column (40 °C), UV and refractive index detectors, and 5 mM H_2_SO_4_ as eluent (0.6 mL/min and 20 µL injection volume). The chromatogram of ammonium contained a constant peak of HEPES buffer interference at a similar elution time, leading to a baseline increase of the ammonium result and a higher variability. Since the buffer concentration should remain relatively constant though, the absolute change in measured ammonium is still representative for the quantities metabolised by the biomass.

### 2.6. Data processing

All images were processed in Fiji (Schindelin et al. (2012)) and background-corrected using a negative control without fluorescent probe or dye. For particle size analysis and shape descriptors, the ‘Extended Particle Analyser’ of the BioVoxxel toolbox was applied (Brocher (2023)). Particle uniformity was assessed as described by Roosen et al. (2015).

Statistical analysis, curve fitting and kernel density estimation (KDE) were performed using SciPy 1.8.0 (Virtanen et al. (2020)) (Python 3.9). Growth rates were fitted as described by D’Huys et al. (2011). All error bars in linear coordinates indicate *µ* ± *σ* with *µ* the sample mean and *σ*^2^ the variance of three biological replicates unless specified otherwise. For logarithmic scales (i.e., for cell and copy numbers), the error bars show the parameters of the lognormal distribution (Magrab (2022)), with *µ* and *σ*^2^ the mean and variance of the variable’s natural logarithm, as follows:

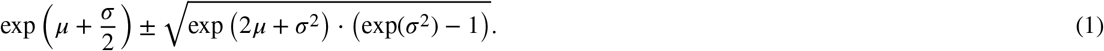

Flow cytometric data analysis and plotting was performed using the FlowCal package in Python (Castillo-Hair et al. (2016)).

## 3. Results

### 3.1 Viable entrapment

To ensure axenic cell entrapment, UV sterilisation of the alginate powder before dissolution was investigated. Sterility was obtained for an alginate load below 3.8 mg/cm^2^ in petri dish experiments, with two UV treatments of 30 min, and intermittent shaking of the powder (supplementary Figure S3). The effect of inhomogeneous distribution on the sterilisation surface was minimised by lowering the load to approximately 1.5 mg/cm^2^ for axenic entrapment experiments. The inoculated alginate beads had an area-equivalent diameter of around 1 mm, with a uniformity ratio of 1.23 (supplementary Figure S4). A worst-case scenario of entrapped activated sludge was monitored to assess the viability and possibly oxygen transfer limitations after encapsulation. It was found that anaerobic processes (reflected by lactic acid production) could be completely suppressed by sufficient aeration (supplementary Figure S5).

The behaviour of all N-cycle bacteria in entrapped conditions was checked individually and compared to a planktonic reference. For planktonic (axenic) cultures, growth rates of 0.88 ± 0.09 d^−1^, 1.12 ± 0.04 d^−1^, and 8.64 ± 0.62 d^−1^ (*µ* ± *6* for three replicates) were observed for AOB, NOB and DEN respectively (see supplementary Section S.2.1). For both entrapped and suspended (planktonic) NOB, the nitrite concentrations are shown in Figure 4. More details (including the results of AOB and DEN) can be found in the supplementary information Section S.2.2, Figures S9-S12. In summary, the uptake of metabolites was not limited by the presence of the alginate matrix. In some cases, it was even slightly faster than the planktonic reference. Considerable amounts of biomass also started to grow in the suspension of reactors containing beads, as observed by optical density and flow cytometric assays on the supernatant. In single-species experiments, this amount of biomass was at least ten times lower than the suspended reference reactor during the first phase of growth. In a coculture reactor run, the combined growth rate of SYTO9 positive, mScarlet-I negative organisms in the supernatant (representing nitrifiers and DEN not expressing (detectable) fluorescent protein) was observed to be 0.55 ± 0.13 d^−1^. An example of such flow cytometric assays over time is provided in supplementary Figure S14. It was also found that the mScarlet-I positive and negative populations show distinct differences in forward and side scatter profiles.

**Figure 4:**
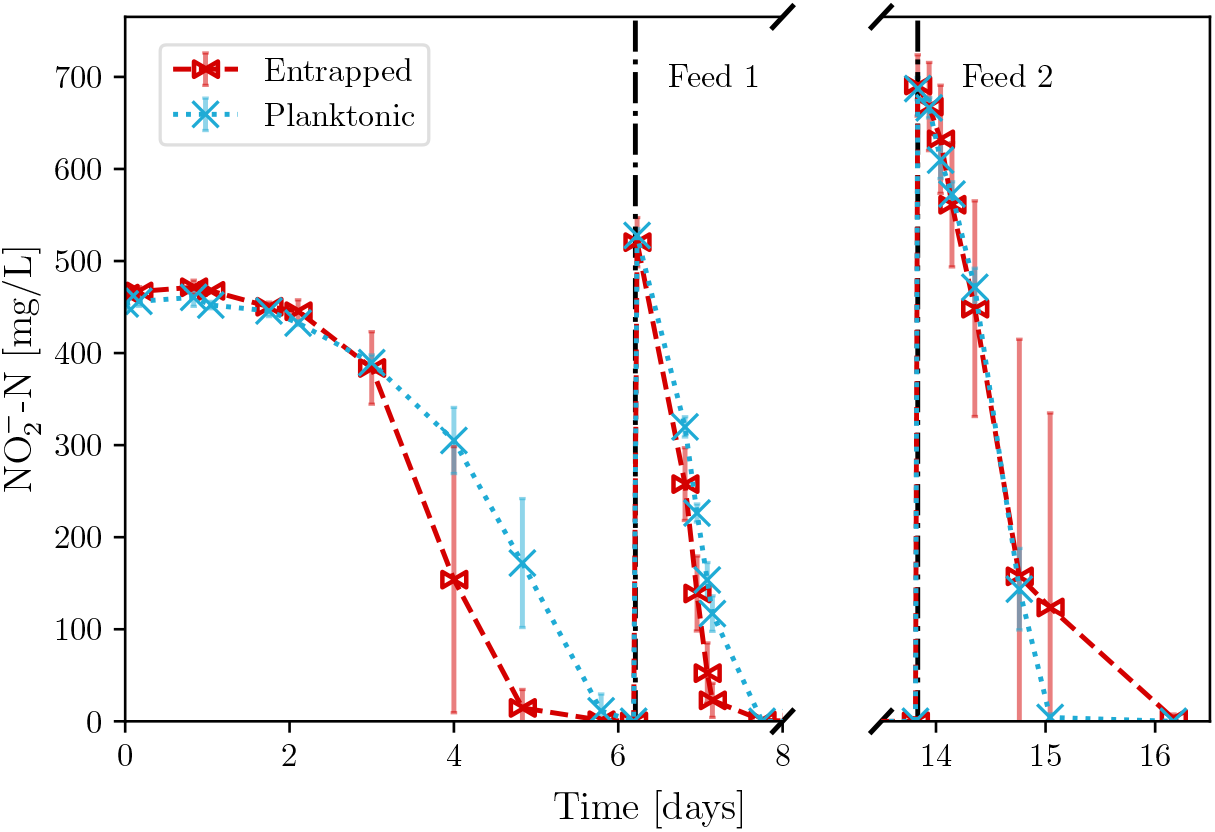
Nitrite concentrations for entrapped and suspended NOB. The uptake of metabolites was consistent for both conditions, and slightly faster (with higher variability) in the first phases of the entrapped case.

### 3.2. Coculture timelapse

In a coculture experiment with all three bacteria, the full nitrification-denitrification cycle was observed during multiple oxic-anoxic cycles, as depicted in Figure 5 for entrapped biomass. The nitrate and nitrite concentrations over time clearly show the nitrification and denitrification activity. Nitrite accumulation occurs in all conditions, with distinctly reduced peaks in the second cycle.

**Figure 5:**
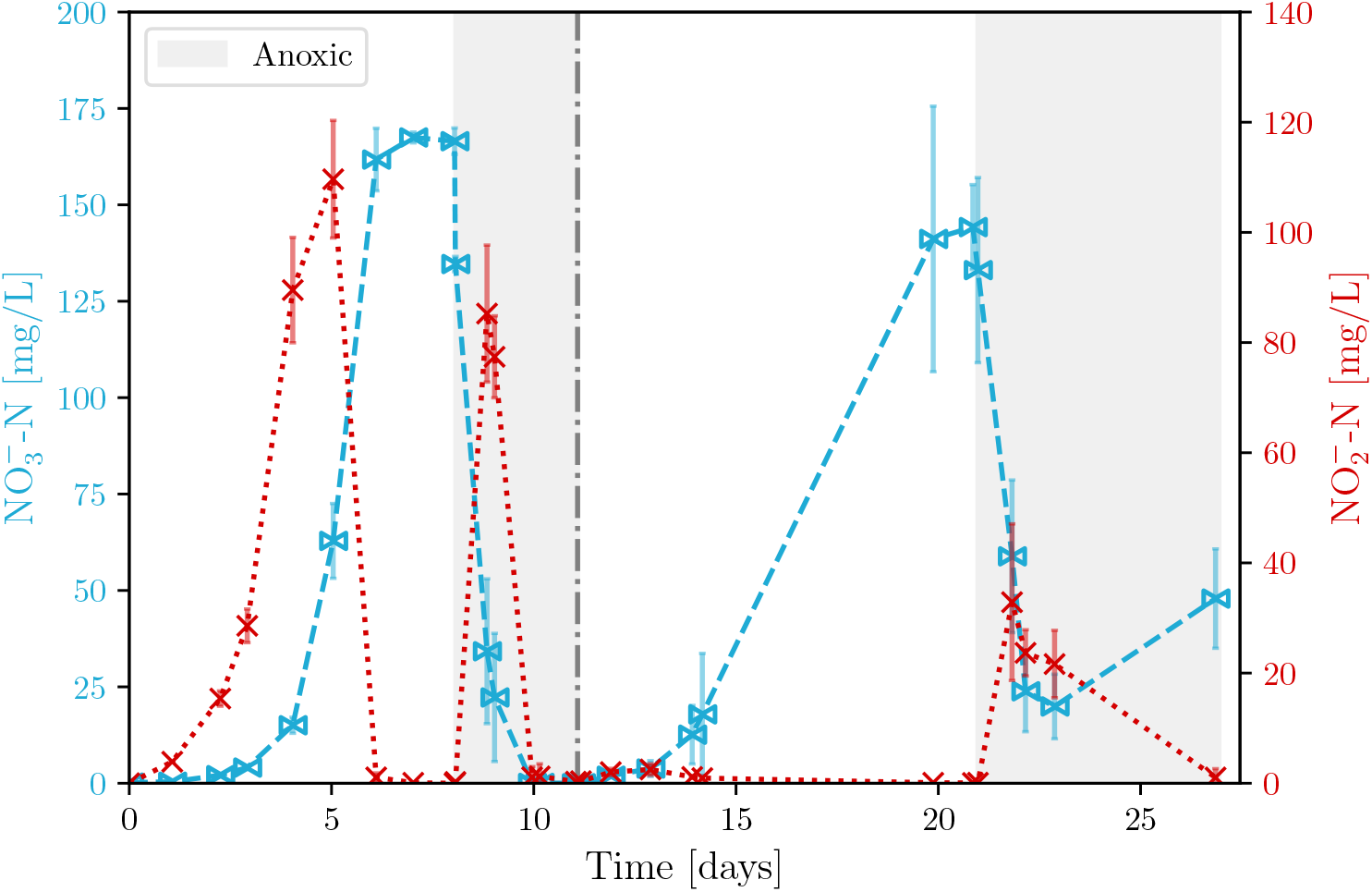
Nitrite (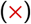) and nitrate (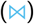) concentrations of the coculture in entrapped conditions for four biological replicates. The nitrite maxima show the intermediate steps in both nitrification and denitrification reactions. After around 12 days, the supernatant was removed and medium was refreshed, as indicated by the dash-dotted line.

To monitor the growth of the entrapped coculture in detail during the first oxic-anoxic cycle, qPCR and cryosectioning-FISH were applied. Figure 6 shows the copy number of the corresponding genes over time. Here, a clear exponential growth of AOB and NOB is observed, with a relatively minor 2-4 fold increase in the gene corresponding to DEN. Note that all three genes show a maximum, reached in the order DEN-AOB-NOB. After the anoxic regime, AOB remained constant, and NOB and DEN decreased.

**Figure 6:**
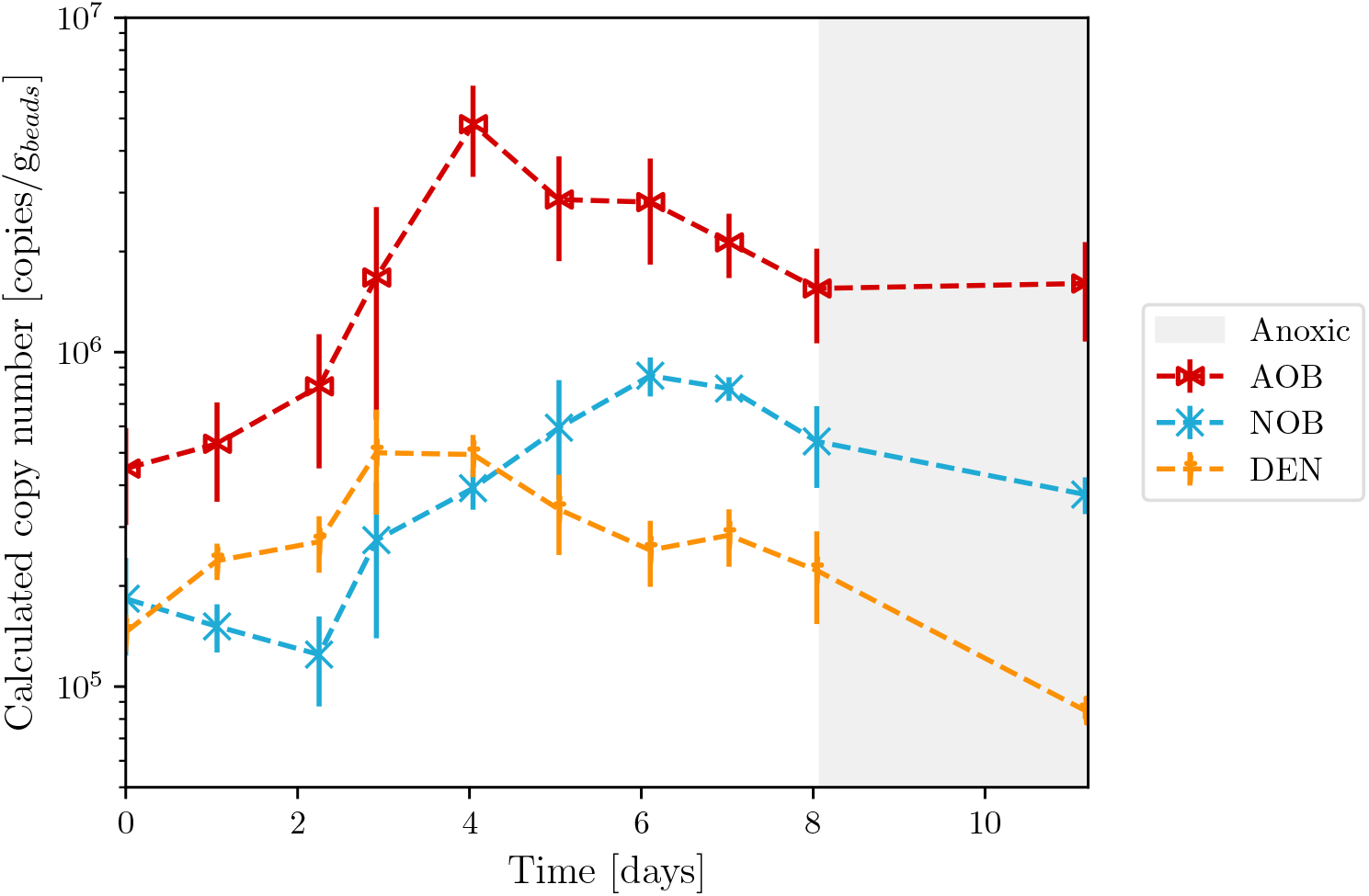
qPCR results for four biological replicates of dissolved alginate beads during the first oxic-anoxic cycle of the entrapped coculture experiment. Copy numbers are expressed per gram of dissolved beads before dissolution and DNA extraction. All curves indicate growth and a maximum in copy number.

Microscopic images of cryosectioned-FISH samples also confirm the colony growth inside the bead (supplementary Figure S15). It should be noted that agarose immobilisation (see Section 2.4) was indispensable to retain the biomass during the FISH procedure. Size distributions of all bacterial colonies over time (based on the EUBMIX-FAM signal) can be found in supplementary Figure S16 and lead to the same observation. After the first aerobic phase, colonies reached an average size of about 4 µm. While the microcolonies are distributed homogeneously throughout the polymer matrix, FISH analysis showed that the species themselves are not. For example, not all locations that contain AOB, also contain NOB or DEN (supplementary Figure S17). In fact, most observations likely correspond to only one of the three species. There was also no observed dependence of colony formation on the radial position inside the bead.

To get an idea of what the entrapped biomass looks like inside the alginate, Scanning Electron Microscope (SEM) micrographs of the sections were acquired as well, as depicted in Figure 7. The scale of the visible colonies is consistent with the ones observed with standard light microscopy. In addition, SEM images more distinctly show the ‘packing’ of the individual bacterial cells in the polymer matrix. Some colonies remain entrapped after sectioning, while others have burst, (partly) releasing the individual cells.

**Figure 7:**
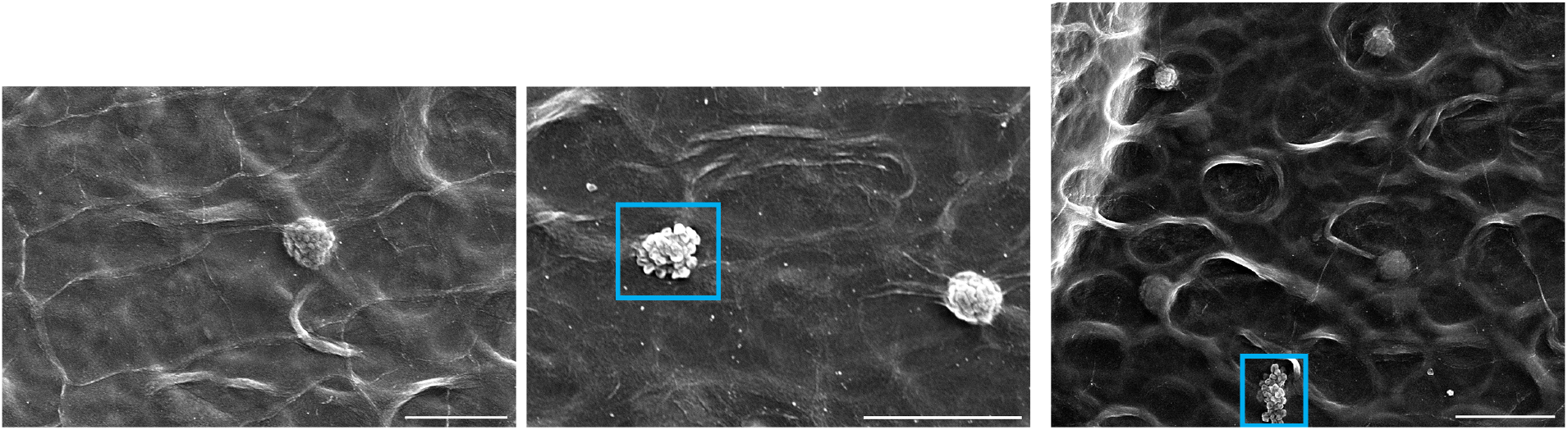
SEM micrographs of a cryosectioned bead after 8 days of incubation. Scale bars indicate 10 µm. Some colonies are tightly packed in the alginate matrix, while others are ‘released’ (cyan rectangles) after sectioning.

### 3.3. Colony release and flocculation

After alginate dissolution by switching to phosphate-buffered medium, the developed colonies were successfully released into suspension. Figure 8 shows the persistence of (N-cycle) metabolic activity after release. The corresponding ammonium concentrations are provided in supplementary Section S.2.4, along with similar metabolite profiles of a reference coculture without initial entrapment. While activity was maintained, the size of the released microcolonies did not change over time. After 9 days of suspended incubation, their average particle size remained around 4 µm (see supplementary Figure S21). An example of typical microcolonies that were observed is provided in Figure 9. From these images, it is clear that no mScarlet-I positive DEN were present in the predominantly spherical structures, suggesting that only nitrifiers are present. The SEM micrographs also indicate that they are tightly packed. These colonies were not observed in planktonic conditions, in contrast to prevalent *Azoarcus* clusters. Figure 10 shows a sample of the released colonies, but subjected to the FISH procedure. In that case, strong clusters were observed alongside individual bacteria. The true content of the bioreactor was therefore not only altered on the scale of the bacteria (due to fixation), but also on a microcolony level.

**Figure 8:**
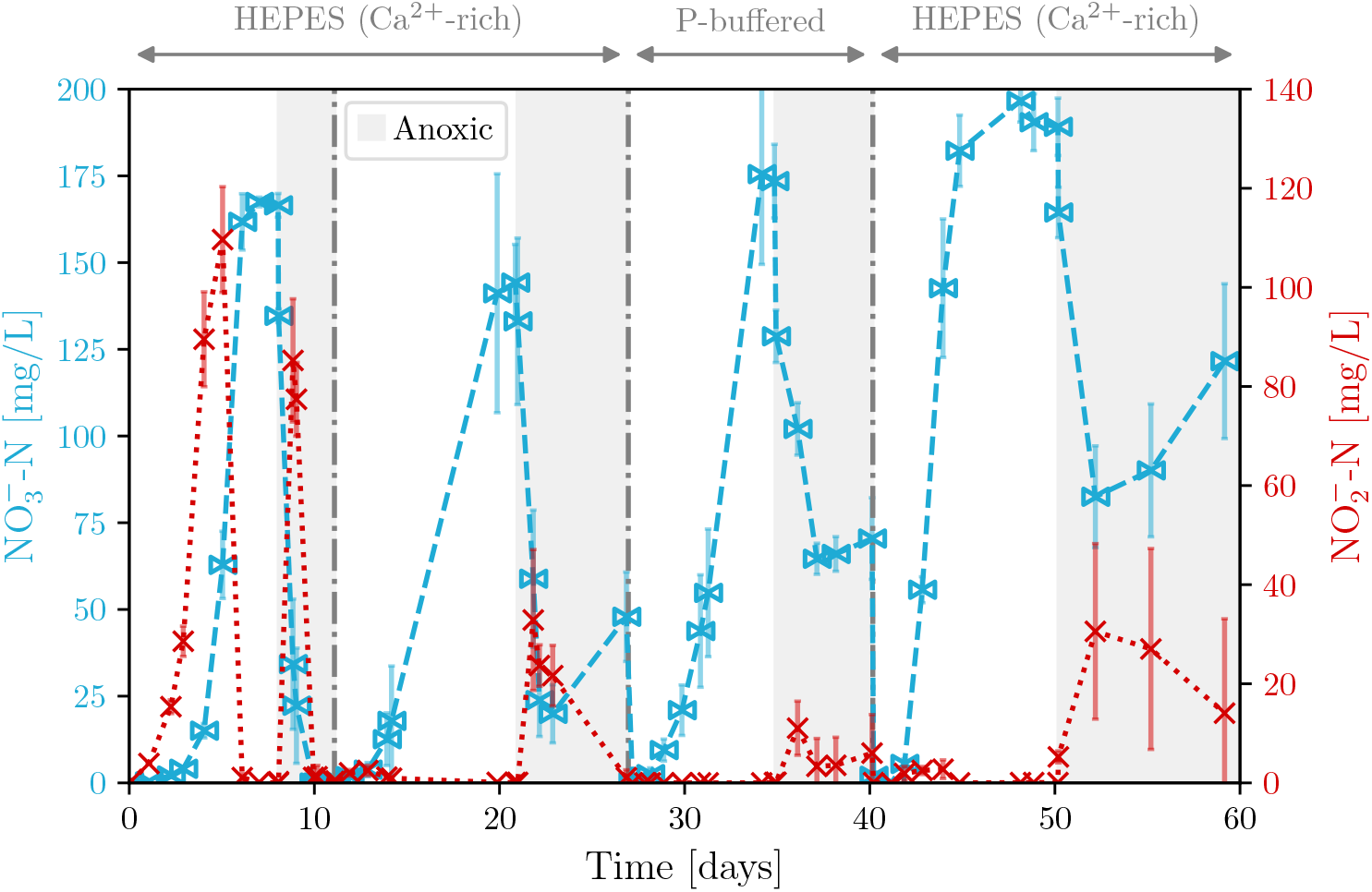
Nitrite 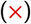 and nitrate 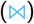 concentrations of the coculture in entrapped conditions for four biological replicates. The supernatant was removed and medium was refreshed at every dash-dotted line. After 27 days, the beads were released by switching to phosphate-buffered medium, and after 40 days the biomass was resuspended again in Ca^2+^ rich medium. The first two cycles correspond to Figure 5.

**Figure 9:**
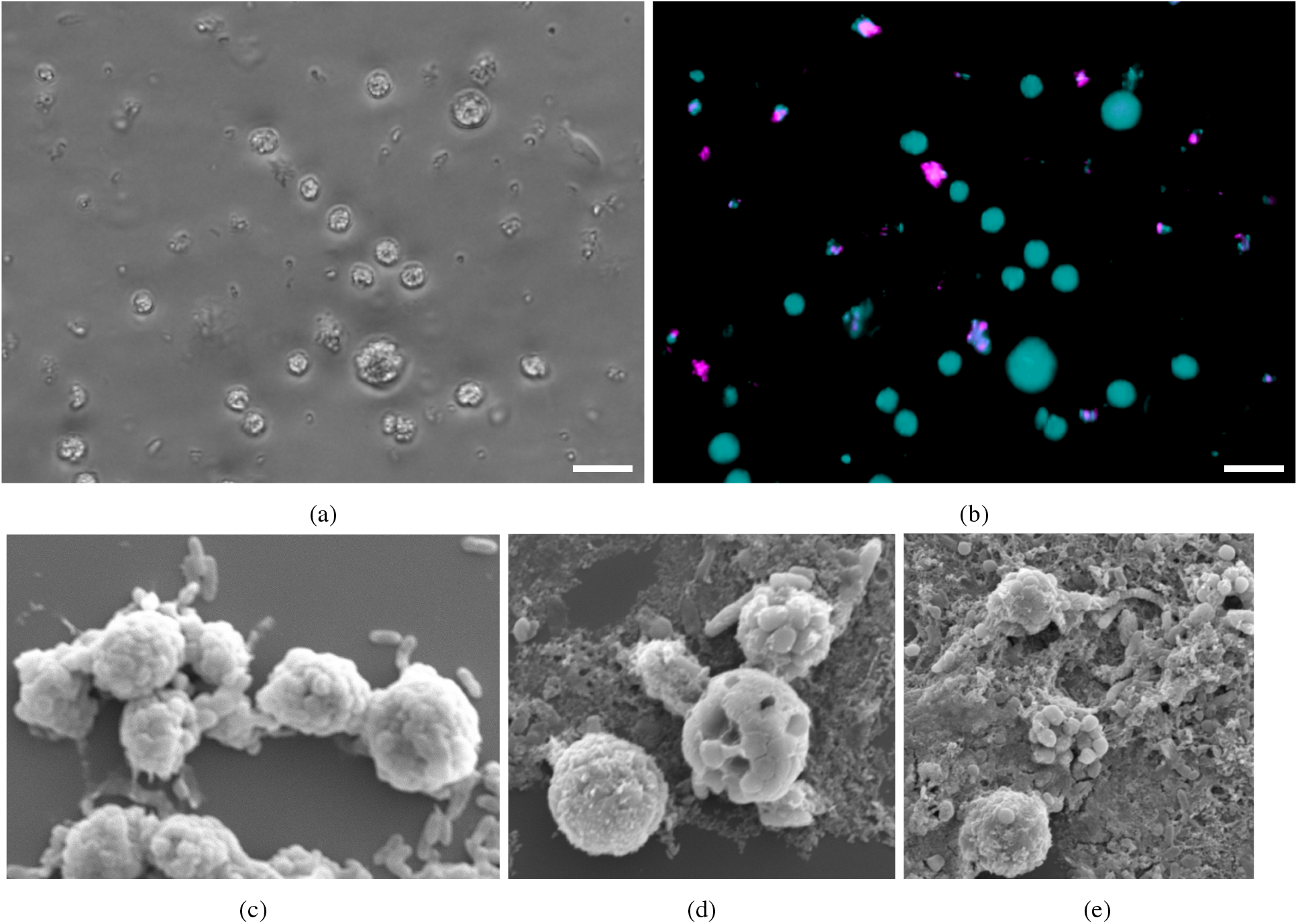
Microscopic images of bacterial colonies in suspension 4 days after alginate bead dissolution. (a) shows a phase-contrast image, with (b) the corresponding fluorescent signals of SYTO9 (all bacteria) in cyan and mScarlet-I (denitrifier) in magenta. (c-d) depict SEM micrographs of the clusters. Scale bars indicate (a,b) 10 µm and (c-e) 5 µm.

**Figure 10:**
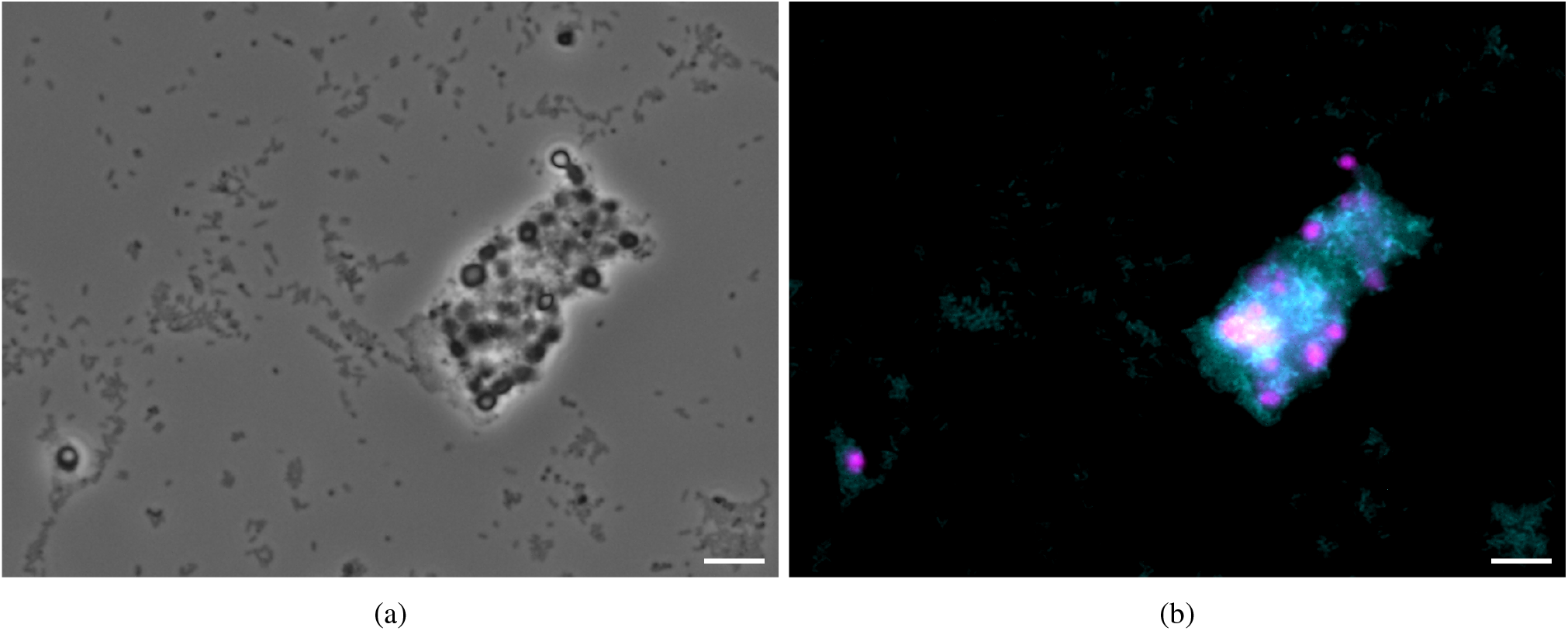
FISH images of bacterial colonies in suspension after alginate solubilisation. Clustering is clearly observed in the phase-contrast image (a) after the FISH procedure, which does not reflect the actual aggregation condition as observed in suspended images (cf. Figure 9). Scale bars indicate 10 µm. (b) shows the overlap of EUBMIX-FAM in cyan and NIT3-Cy3 in magenta (containing mScarlet-I from DEN as well).

To induce the formation of macroclusters in suspension, the biomass was centrifuged at 3000 g for 10 min, and resuspended again in calcium-rich, HEPES-buffered medium. Microscopic images of the suspension in a well plate showed considerable aggregation as shown in Figure 11. The result was followed up for another 10 days, and similar floc-like structures remained present in aerobic and stirred conditions (see supplementary Section S.2.6). Individual colonies and bacteria were also observed alongside the larger structures throughout the complete run. While centrifugation and resuspension agglomerated the existing (dense) structures (cf. Figure 11), new (characteristic) *Azoarcus* clusters were also forming during the aerated phase after the medium switch, and these contained nitrifier colonies in a looser (more transparent) matrix (for example supplementary Figure S22).

**Figure 11:**
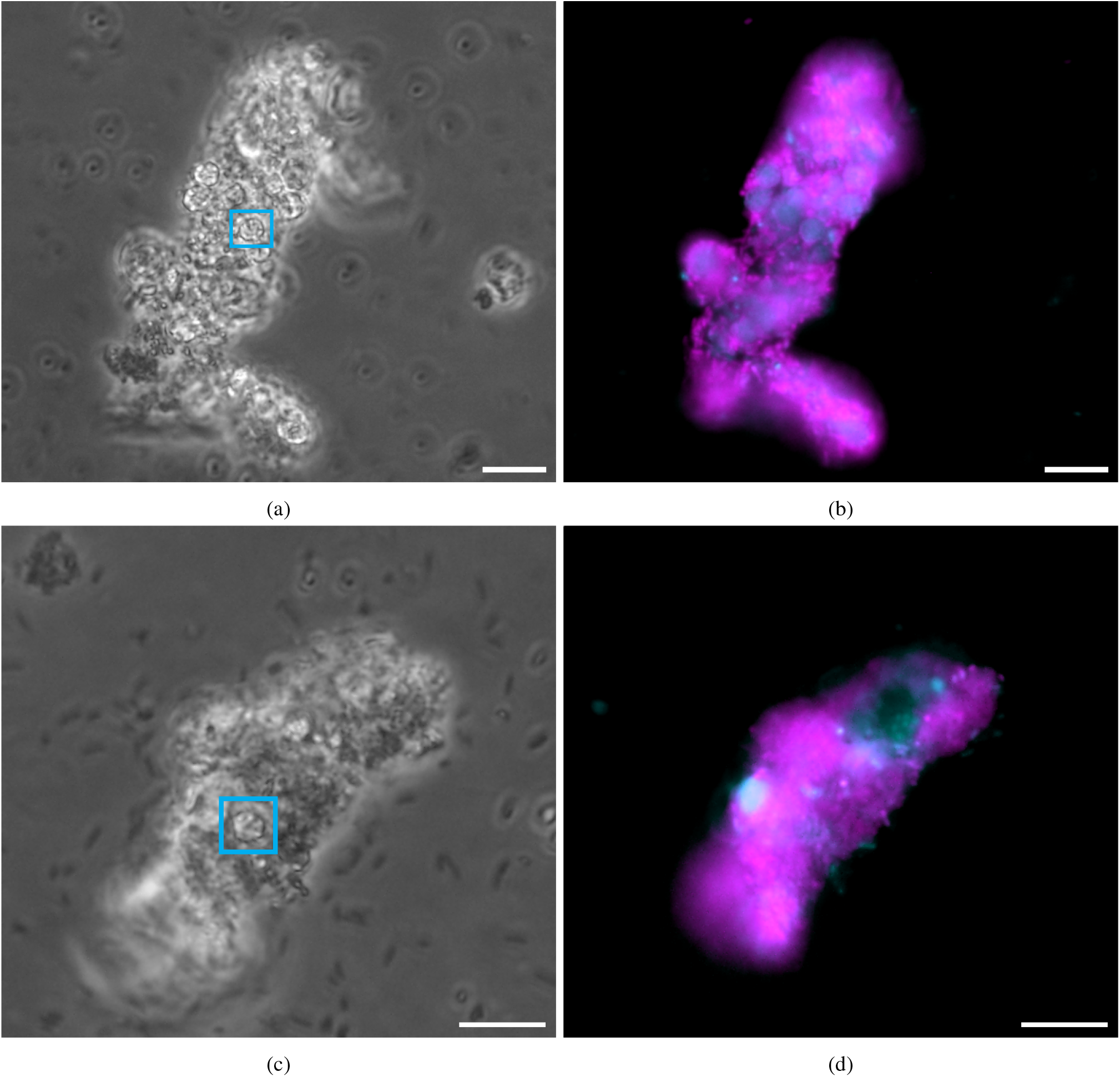
Microscopic images of suspended macroclusters after centrifugation and resuspension in Ca^2+^-rich HEPES-buffered medium. (a,c) show phase-contrast images, with (b,d) the corresponding fluorescent overlapped signals of SYTO9 in cyan and mScarlet-I in magenta. Cyan rectangles indicate examples of nitrifier colonies similar to Figure 9. Scale bars indicate 10 µm.

To investigate the effect of centrifugation, the size of *Azoarcus* clusters was determined before and after centrifugation (and resuspension in calcium-rich medium) in a reference experiment with the planktonic coculture. From image analysis, it was found that centrifugation led to temporal compaction and therefore larger clusters of mScarlet-I positive cells. These aggregates did fall apart in less than two days under aerobic and stirred conditions. More details can be found in the particle sizes and characteristics depicted in supplementary Figures S27-S29.

## 4. Discussion

### 4.1 Viable entrapment

Since standard sterilisation techniques such as autoclaving (Lorson et al. (2020)) and gamma irradiation (Chang et al. (2022)), alter the physicochemical properties of the polymer, UV sterilisation of sodium alginate powder was investigated. While Lorson et al. (2020) advised against the use of UV irradiation due to an ineffective sterilisation outcome, Yu et al. (2017) showed good results for two UV cycles of 25 min. As shown in this work, the outcome does not only depend on the sterilisation time, but especially on the powder load on the sterilisation surface and its distribution. For this reason, care should be taken to minimise inhomogeneities (for example by intermittent redistribution) and the maximum alginate load should be reevaluated when applying a different UV lamp or setup. In any case, alginate sterilisation is indispensable when working with (slow growing) pure cultures, as the contamination level in this (natural) product is unacceptable for this purpose.

Although beads were generated with considerable sizes for oxygen diffusion limitations (∼1 mm), no metabolic issues were observed. Furthermore, there was no dependence of the growth/survival of microcolonies on their radial location inside the calcium alginate beads, suggesting that no limitations were present in the described experiments. Note that the diffusivity of the nitrogen species themselves inside the polymer matrix were not determined, but can be measured according to Landreau et al. (2021). A more in-depth study on this topic (including modelling) was already performed by Wang et al. (2022) for *Nitrosomonas*. Note that not only carrier size, but also porosity, viscosity and many gelation parameters (time, pH, concentration) affect substrate limitations, as for example reviewed by Cassidy et al. (1996). During entrapped pure culture tests with *Nitrobacter*, an accidental leak in the air line showed that the uptake of nitrite became linear (and slower) at low air flow rates (see supplementary Figure S13), again indicating the absence of rate-limiting steps in normal operating conditions. Individual entrapment exhibited a slightly faster metabolite uptake in the initial phase compared to a planktonic reference for AOB and NOB, but not DEN.

### 4.2. Coculture timelapse

The coculture was successfully monitored over time using qPCR, metabolite analysis and microscopy. All tools showed the growth of all three species. Especially in the first growth phase, nitritation is clearly observed during the lag phase of *Nitrobacter* (NOB), before full nitrification to nitrate sets in. In the anoxic phases, nitrite accumulation is observed as well. This phenomenon depends on the applied carbon source and COD to 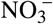 ratio, as described by Ge et al. (2012). In this work, however, the biomass concentration (and environmental adaptation) may be more relevant to explain the decreasing nitrite peaks.

Concerning the gene copy number, the bacteria reach their maximum inside the beads in a logical relative order according to their growth rate and metabolite availability. NOB follows AOB, due to the former’s dependence on nitritation. Both AOB and NOB also decrease in copy number after their main metabolites (ammonium and nitrite respectively) have been depleted. The amoA gene of *N. europaea* (AOB) reaches a maximum copy number that is 5-6 times higher than the nxrB gene of *N. winogradskyi* (NOB), which was not observed with respect to cell counts in pure culture growth. While the number of genomic copies in each cell can have an influence, this is identical for the genes targeted here (Hommes et al. (1998); Starkenburg et al. (2006)). Pérez et al. (2014) did show, however, that *N. europaea* benefits more from a coculture than *N. winogradskyi*, with the ammonium oxidising bacterium taking up four times more biomass in a coculture chemostat. In any case, the results show that TaqMan qPCR on the entrapped coculture is a feasible and reproducible way to quantify biomass and monitor the growth of individual species. Other techniques to obtain a proxy for biomass content such as total protein quantification on the dissolved beads (see for example Gutenberger et al. (2024)), are more relevant for larger communities and were therefore not applied here.

After the first aerobic growth phase, the entrapped colonies reached their maximum size of around 4 µm. This was confirmed by micrographs of released microcolonies, and means that in the intermediate phases before release, not much changed with respect to bead content. The FISH analysis did show that the entrapped biomass remains active (targeting the 16S rRNA), and metabolically, the coculture is able to retain its function after washing and supernatant replenishment. *Azoarcus* also colonises the supernatant in this intermediate stage, indicating that the beads (and possibly some biofilm formation on the vessel wall) still serve as a viable inoculum for the denitrifier as well, even though qPCR and microscopic images indicate its presence inside the beads was limited. The reason for the lower amount of DEN in the coculture bead might be due to both (or the combined effect of) its motility and limited/suppressed feed (200 mg Ac^−^/L in coculture conditions). After all, in pure culture experiments, the whole bead was colonised by *Azoarcus* as well.

Regardless of the inoculum composition, the microcolony formation inside the beads was always homogeneous. However, FISH analysis confirmed that not all species are present in each colony. It might therefore not be advantageous to occupy the same (rather limited) space in the alginate matrix. For example, colonies containing both AOB and NOB do not have a metabolic advantage over colonies with only AOB or NOB, because nitrite diffusion is not limiting within the bead. In addition, it is also possible that not all points of inoculation lead to complete colony formation as a result of competition. However, the resolution of the microscopic images is not high enough to determine the number of initial inoculation points per section area. Nevertheless, an important observation from the data shown here is that the studied nitrifiers are in fact compacted into a strong cluster, possibly due to the steric hindrance inside the cavities.

### 4.3. Colony release and flocculation

After a switch to phosphate-buffered medium, the precipitation of calcium phosphate quickly led to the reversal of the ionotropic alginate gelation and the release of entrapped biomass. Especially the spherical nitrifier clusters were immediately apparent in suspension and were consistently observed for 24 days. The SEM micrographs confirmed the distributed growth of the species, because the constituents of the clusters mostly had the same morphology (*Azoarcus* is rod-shaped, unlike the nitrifiers). Important to stress here is that macroclusters were not observed for samples in well plates, while FISH-stained samples did show artefactual aggregation. This observation highlights the impact of sample preparation when considering interspecies localisation: the FISH procedure itself consistently brought the individual building blocks together, which did not reflect the actual reactor content. In an attempt to achieve the same level of attachment in suspension, the reactor content was centrifuged and resuspended again in calcium-rich medium. This medium change led to the combination of nitrifier colonies with the typical suspended *Azoarcus* flocs into a *de novo* floc containing all species. While the macroaggregation confirms the cation bridging theory postulated by Sobeck and Higgins (Sobeck and Higgins (2002)), the results of batch experiments as shown here are not ideal for this purpose. Sobeck and Higgins even recommend at least 2-3 sludge retention times (SRT) in sequencing batch reactor (SBR) operation before drawing conclusions (Sobeck and Higgins (2002)). Indeed, the effect of centrifugation should not be underestimated as well. In a planktonic reference, *Azoarcus* cluster size seemed to be only temporarily affected: the initial compaction after centrifugation was reversed in less than 2 days. Nevertheless, the centrifugation and cation bridging cannot be fully decoupled in the results of these experiments. Therefore, full (automated) SBR or continuous operation is advised for future research, to be able to switch the ionic content of the reactor without interfering with the formed aggregates by centrifugation.

## 5. Conclusion

This work aimed to establish a bottom-up approach to study bioflocculation. The applied N-cycle consortium could be reproducibly grown in coculture conditions, and was monitored in oxic/anoxic cycles using microscopy and metabolite measurements. To force the bacteria together, the biomass was first entrapped in calcium alginate beads, and released again after two complete cycles. The following conclusions were drawn from the results of these experiments.

- The suitability of alginate entrapment as cell retention system proved useful in this type of experiments, since no detrimental effect on the metabolic activity of the bacteria was observed.
- An EPS producer is a solid starting point, because *Azoarcus communis* did not even need entrapment to form flocs, and attachment to this bacterial matrix was possible. With this in mind, the extension of the studied consortium with extra functionality, such as phosphate accumulating organisms (PAO), seems feasible and could provide more insights into their role in flocculation and granulation.
- Fluorescent in-situ hybridisation (FISH) was not suitable to provide clustering information since the procedure itself changed the apparent aggregation state of the reactor content.
- qPCR after bead dissolution is a reproducible tool to quantify entrapped biomass and can be extended to follow up any genes or species of interest.
- True (automated) sequencing batch reactor (SBR) operation is advised when studying flocculation to avoid centrifugation for medium exchange. In this way, the impact of calcium can be fully decoupled from (temporary) centrifugation effects.
- Strong, spherical nitrifier colonies were identified during and after entrapment. These compact aggregates were active, stable, and of constant size after release into suspension. Since these structures were not observed in planktonic cultures, it would be interesting to investigate the source of this phenotype using for example Raman spectroscopy and omics techniques. After all, they have been observed to form strong microcolonies in activated sludge as well.

## Supplementary materials

Supplementary information can be found in the online version of the article in supplementaryinfo.pdf

## Acknowledgements

The authors would like to thank Prof. Kristel Bernaerts for her helpful input in all stages of the project; Prof. Karoline Faust and Prof. Maarten Roeffaers for the use of the flow cytometer and SEM, respectively; Rik Nuyts for his training on confocal microscopy; Greet Van de Velde for her help with the IC analysis; Sulieman Minhas for his tips and techniques to obtain good sections with the cryostat; Pieter De Wever for his help with the encapsulator; Dries Grauwels for sharing the procedure to generate qPCR standard template.

## CRediT authorship contribution statement

**Laurens Parret:** Conceptualisation, Methodology, Investigation, Writing - Original draft preparation, Visualisation. **Kenneth Simoens:** Conceptualisation, Methodology. **Benjamin Horemans:** Conceptualisation, Methodology. **Jo De Vrieze:** Conceptualisation, Writing - Review & Editing, Supervision, Funding acquisition. **Ilse Smets:** Conceptualisation, Writing - Review & Editing, Supervision, Funding acquisition.

