## Supplementary Information for "Establishing a co-culture aggregate of N-cycle bacteria to elucidate flocculation in biological wastewater treatment"

### S.1 Methods

**Table S1**: Coculture medium composition (based on DSMZ medium 1583)

| Compound | Concentration [mg/L] | HEPES medium | P-buffered dissolution medium | Preculture AOB ($\alpha$) NOB ($\beta$) |
|---|---|---|---|---|
| $NH_4Cl$ | 200 mg-N | × | × | $\alpha$ |
| $NaNO_2$ | 200 mg-N | | | $\beta$ |
| $CaCO_3$ | 70 | | | × |
| Sodium acetate | 0-200 mg-Ac | | × | |
| Calcium acetate | 0-200 mg-Ac | × | | |
| HEPES | 6000 | × | | |
| $KH_2PO_4$ (minimal P) | 5.4 | × | | |
| $CaCl \cdot 2\,H_2O$ | 147 | × | | |
| $Na_2HPO_4$ (P-buffer) | 1420 | | × | × |
| $KH_2PO_4$ (P-buffer) | 240 | | × | × |
| Yeast extract (Difco) | 200 | × | × | |
| Fixed compounds | | × | × | × |
| NaCl | 584 | | | |
| KCl | 74 | | | |
| $MgSO_4 \cdot 7\,H_2O$ | 49 | | | |
| Trace element solution | 1 mL | | | |
| Cresol red solution | 2 mL | | | |
| pH | 7.8 | | | |
| Trace element solution | | | | |
| HCl, 1 M | 25 mL | | | |
| $MnSO_4 \cdot 4\,H_2O$ | 45 | | | |
| $H_3BO_3$ | 49 | | | |
| $ZnSO_4 \cdot 7\,H_2O$ | 43 | | | |
| $(NH_4)_6Mo_7O_{24} \cdot 4\,H_2O$ | 37 | | | |
| $FeSO_4 \cdot 7\,H_2O$ | 973 | | | |
| $CuSO_4 \cdot 5\,H_2O$ | 25 | | | |
| Cresol red solution | | | | |
| Cresol red | 500 | | | |

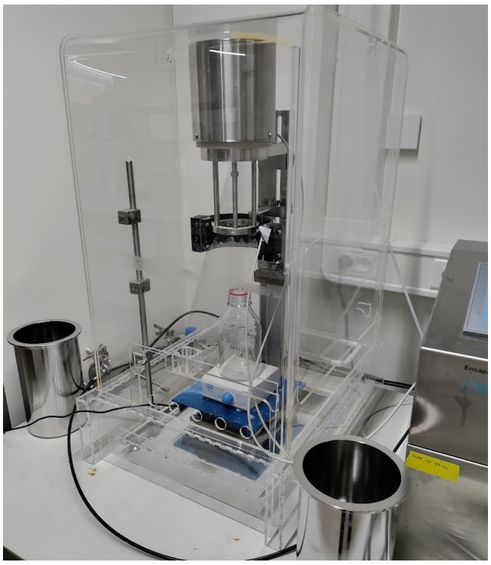

**Fig. S1**: Axenic bead production setup with a plexiglass box around the Nisco encapsulator.

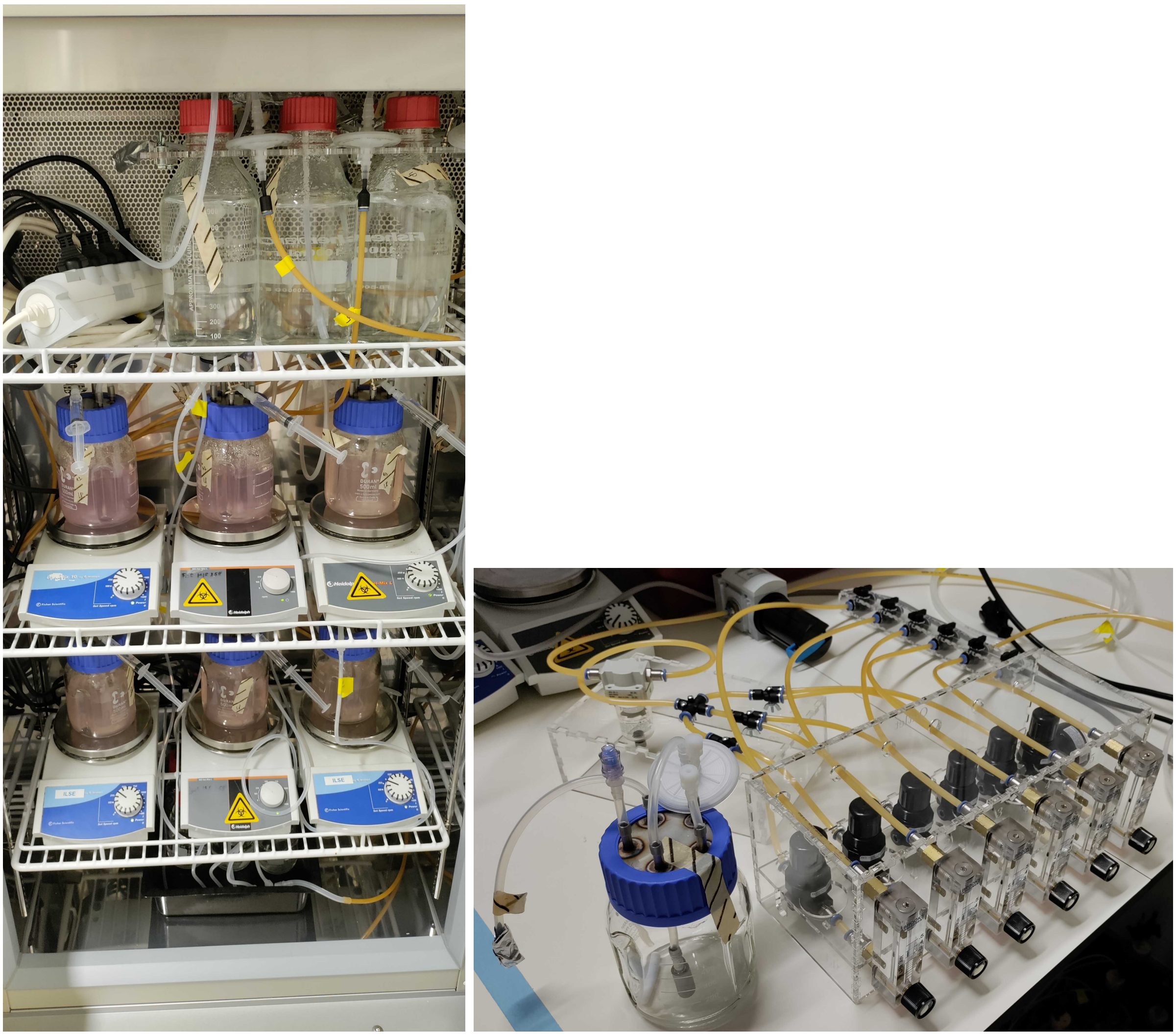

(a) (b)

**Fig. S2**: Pictures of (a) the reactor setup and (b) the aeration module.

#### S.1.1 FISH

**Table S2**: Applied FISH probes along with their sequence and FA concentration. Details on oligonucleotide probes and their references are available at probeBase[1].

| Probe name | Target organism | Sequence | Applied fluorophore(s) | %FA |
|---|---|---|---|---|
| NEU | AOB | CCC CTC TGC TGC ACT CTA | Atto-390, Cy5 | 40 |
| NIT3 | NOB | CCT GTG CTC CAT GCT CCG | Cy3, Cy5 | 40 |
| EUB-MIX (EUB338 + EUB338 II + EUB338 III) | All bacteria | GCT GCC TCC CGT AGG AGT<br>GCA GCC ACC CGT AGG TGT<br>GCT GCC ACC CGT AGG TGT | FAM | (0-50) |

[1]Greuter, D., Loy, A., Horn, M., Rattei, T.: probebase—an online resource for rRNA-targeted oligonucleotide probes and primers: new features 2016. Nucleic Acids Research 44(D1), 586-589 (2015) https://doi.org/10.1093/nar/gkv1232

#### S.1.2 PCR and qPCR

To generate DNA template, the conditions in Table S3 were applied with DNA extracted from pure cultures of each individual strain and their corresponding primers (Table S4) at a final concentration of 300 nM. The PCR product was then separated on a 1.5 wt% agarose gel for 90 min at 90 V, cut out using a scalpel and cleaned up using a Qiagen PCR cleanup kit. The final DNA concentration was determined using a Qubit 1X assay (Invitrogen) and converted into copy numbers to use as standard template. For qPCR, probe and primer concentrations were kept at 200 nM, except for the AOB qPCR probes (primers 900 nM, probe 250 nM according to the mix applied by Orschler et al.[2]. The qPCR conditions are detailed in Table S3). All oligonucleotides developed in this work were designed using NCBI Primer-BLAST[3] with as input the cited sequences in Table S4. The oligonucleotides that were taken from other sources are also provided in Table S4.

**Table S3**: PCR and qPCR conditions.

| | Step | Temperature [°C] | Time |
|---|---|---|---|
| | Activation | 95 | 10 min |
| PCR standard template | 30 cycles | | |
| | Denaturation | 95 | 60 s |
| | Annealing | 55 | 30 s |
| | Extension | 72 | 60 s |
| | Stop cycling | | |
| | Extension | 72 | 10 min |
| | Activation | 95 | 3 min |
| qPCR | 40 cycles | | |
| | Denaturation | 95 | 10 s |
| | Annealing/Extension | 60 | 60 s |

[2]Orschler, L., Agrawal, S., Lackner, S.: Lost in translation: the quest for Nitrosomonas cluster 7- specific amoA primers and TaqMan probes. Microbial Biotechnology 13(6), 2069–2076 (2020) https: //doi.org/10.1111/1751-7915.13627

[3]Ye, J., Coulouris, G., Zaretskaya, I., Cutcutache, I., Rozen, S., Madden, T.L.: Primer-BLAST: A tool to design target-specific primers for polymerase chain reaction. BMC Bioinformatics 13(1) (2012) https://doi.org/10.1186/1471-2105-13-134

**Table S4**: Primers and probes for standard template generation by PCR, and for qPCR analysis.

| | Organism | Gene | Forward primer | Reverse primer | TaqMan probe | Source |
|---|---|---|---|---|---|---|
| PCR standard template | AOB | *amoA* | (amoA-1F) GGGGTTTCTACTGGTGGT | (amoA-2R) CCCCTCKGSAAAGCCTTCTTC | / | [4] |
| | NOB | *nxrA-nxrB* | (nxrA-1F) GCATGGATCCGGTGTGGATCA | (NxrB 1R) CCGTGCTGTTGAYCTCGTTGA | / | [5] |
| | DEN | *nifH* | (nifH-Az-1F) CAAATCCACCACCACCCAGA | (nifH-Az-1R) CCGACGATGGAGGTGTCTTC | / | This work (sequence by [6]) |
| qPCR | AOB | *amoA* | (nerF) GTCCCATGTAATCAGCCATC | (nerR) CACACTACCCCATCAACTTC | 6-FAM (nerTaq) BHQ-1 ATAGAACAGCAGACCGAAGAATCCACCTCCAACCA | [7] |
| | NOB | *nxrB* | (nxrB-2q_F) CGACTATTTCGAGCCGTGGA | (nxrB-2q_R) CGCCTCAATCGTGTCCATGT | JOE (nxrB-2q_Taq) BHQ-1 AACCTGATCACTGCGCCTCTGGCTGACGA | This work (sequence by [5]) |
| | DEN | *nifH* | (nifH-Az-2q-F) TGCAACACGCTGAGATCCG | (nifH-Az-2q-R) ATCGGGGTCGGGATAACGAA | ROX (nifH-Az-2q-Taq) BHQ-2 ACCGTGCCCTGGCTCGCAAGATCGTCGATA | This work (sequence by [6]) |

[4] Rotthauwe, J.H., Witzel, K.P., Liesack, W.: The ammonia monooxygenase structural gene amoA as a functional marker: molecular fine-scale analysis of natural ammonia-oxidizing populations. Applied and Environmental Microbiology 63(12), 4704–4712 (1997) https://doi.org/10.1128/aem. 63.12.4704-4712.1997

[5] Vanparys, B., Spieck, E., Heylen, K., Wittebolle, L., Geets, J., Boon, N., Vos, P.D.: The phylogeny of the genus Nitrobacter based on comparative rep-PCR, 16s rRNA and nitrite oxidoreductase gene sequence analysis. Systematic and Applied Microbiology 30(4), 297–308 (2007) https://doi.org/10. 1016/j.syapm.2006.11.006

[6] Raittz, R.T., Pierri, C.R.D., Maluk, M., Batista, M.B., Carmona, M., Junghare, M., Faoro, H., Cruz, L.M., Battistoni, F., Souza, E., Oliveira Pedrosa, F., Chen, W.-M., Poole, P.S., Dixon, R.A., James, E.K.: Comparative genomics provides insights into the taxonomy of Azoarcus and reveals separate origins of nif genes in the proposed Azoarcus and Aromatoleum genera. Genes 12(1), 71 (2021) https://doi.org/10.3390/genes12010071

[7] Orschler, L., Agrawal, S., Lackner, S.: Lost in translation: the quest for Nitrosomonas cluster 7- specific amoA primers and TaqMan probes. Microbial Biotechnology 13(6), 2069–2076 (2020) https: //doi.org/10.1111/1751-7915.13627

### S.2 Results

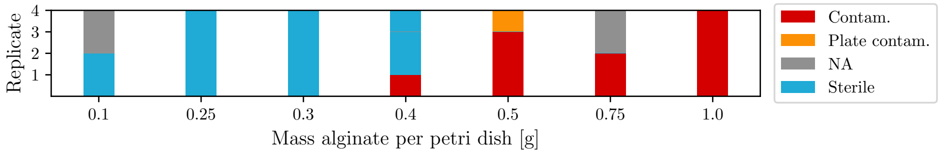


**Fig. S3**: Contamination of alginate powder suspended in Tryptic Soy Broth (TSB) after UV sterilisation. A threshold value for the layer thickness to ensure sterility was determined. Liquid cultures were plated on Tryptic Soy Agar (TSA) to verify any lack of visual contamination.

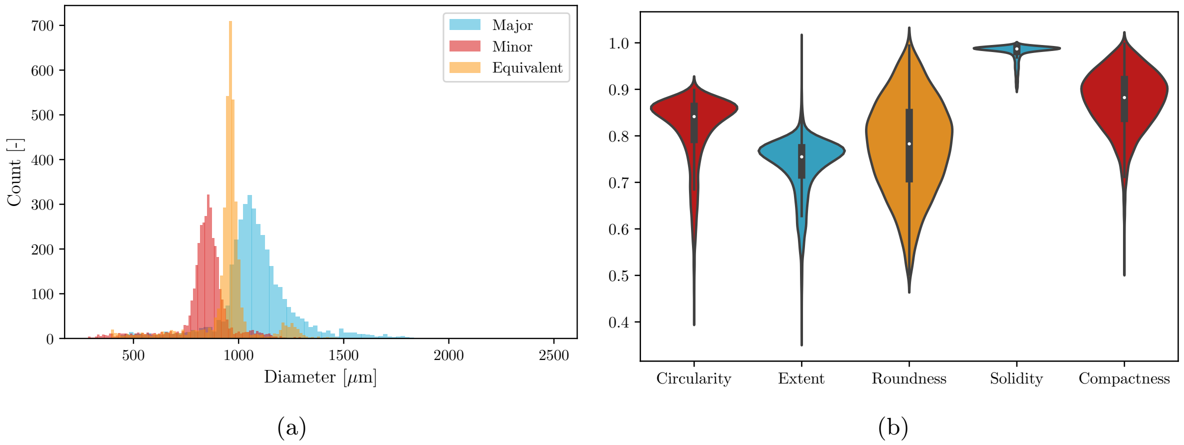


**Fig. S4**: Particle size analysis as determined from brightfield images of crystal violet stained alginate beads. a) Shows the major and minor diameters, as well as the equivalent diameter based on area. b) Indicates violin plots (kernel density estimation) of their shape parameters.

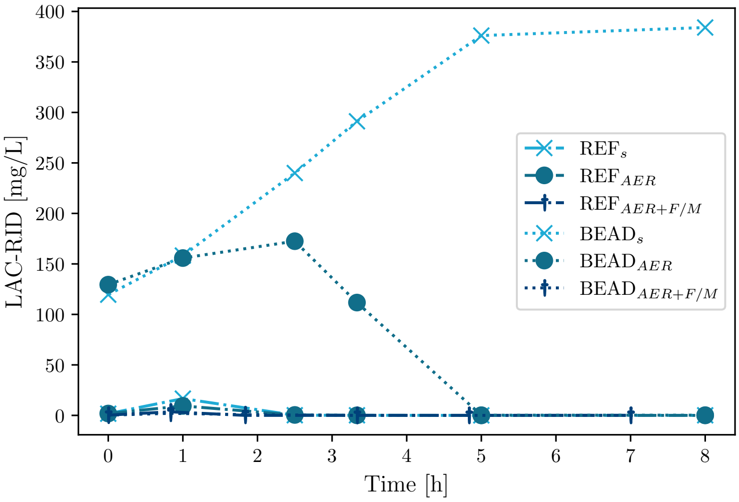


**Fig. S5**: Lactic acid (LAC) concentrations for both suspended (REF) and encapsulated (BEAD) sludge, under stirred (S), aerated (AER), and aerated with constant F/M (food/microorganisms ratio) conditions (AER+F/M). This last variation was necessary to eliminate lactic acid from previous cycles, because at higher feed to entrapped biomass concentrations, there was simply not enough sludge present to metabolise it in time (after LAC production). The anaerobic processes are suppressed with sufficient aeration, indicating no measured effect of oxygen limitations.

#### S.2.1 Growth curves

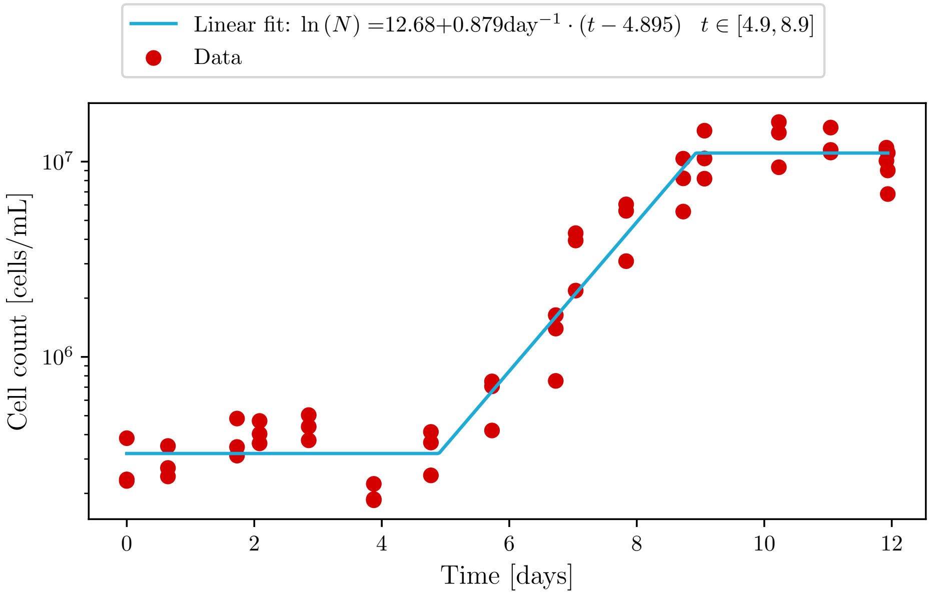


**Fig. S6**: Growth curve of *Nitrosomonas europaea* based on cell numbers (flow cytometry).

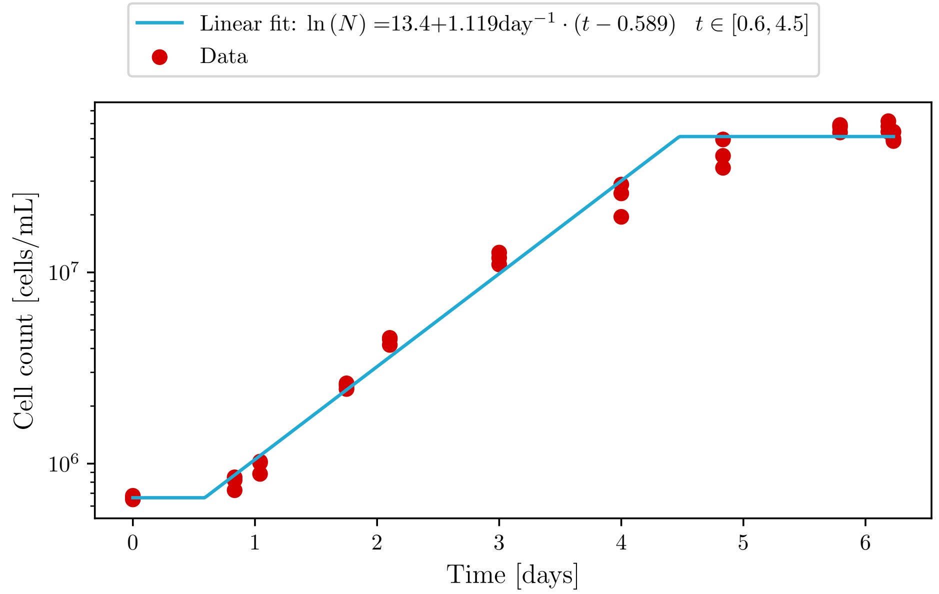


**Fig. S7**: Growth curve of *Nitrobacter winogradskyi* based on cell numbers (flow cytometry).

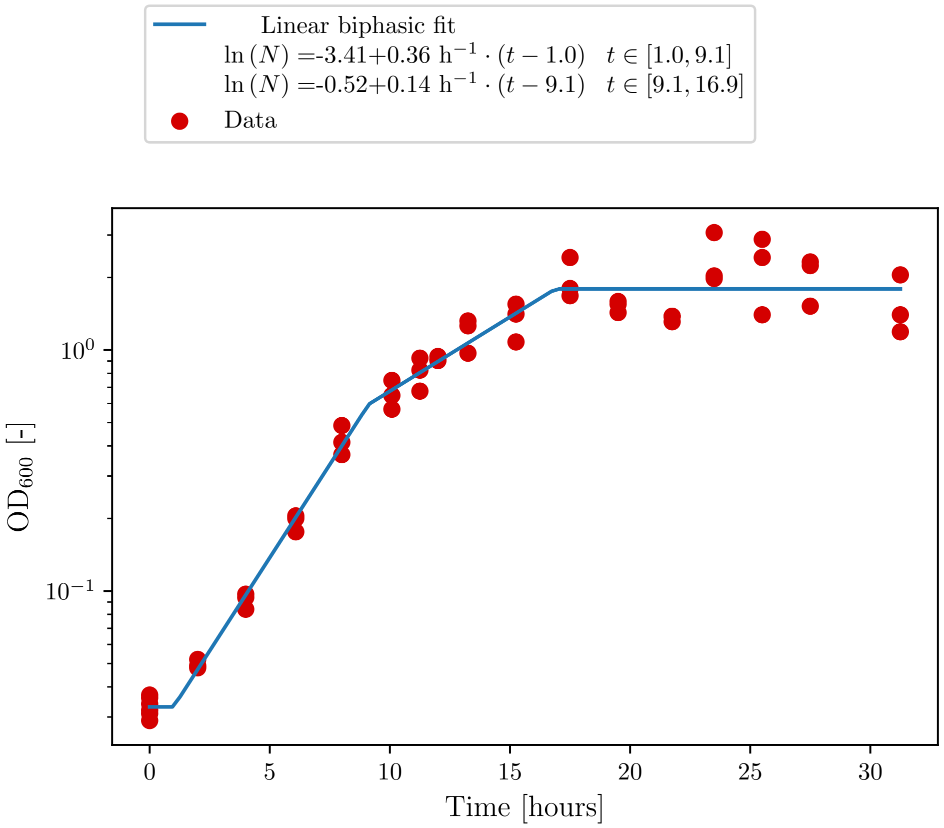


**Fig. S8**: Growth curve of *Azoarcus communis* $\text{Rif}^r$-mSc based on optical density ($OD_{600}$).

#### S.2.2 Axenic entrapment

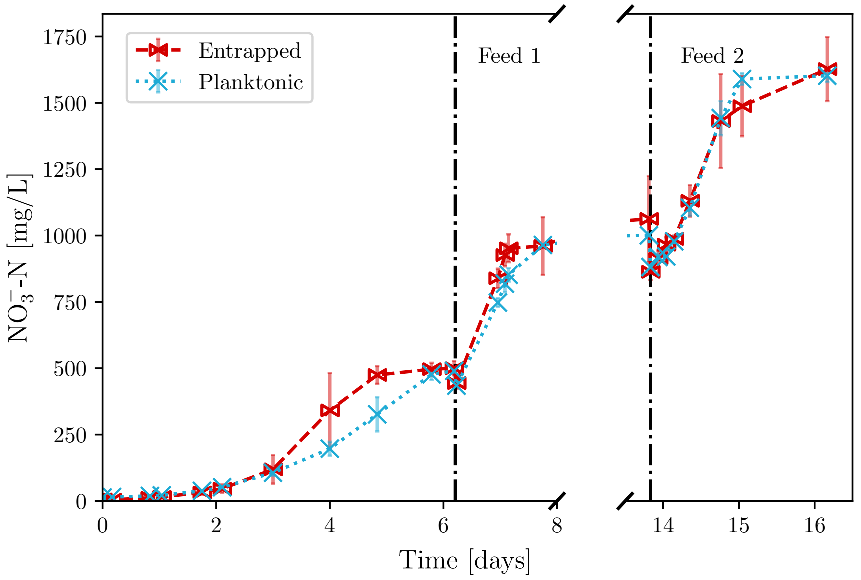


**Fig. S9**: Nitrate concentrations for entrapped and suspended NOB corresponding to Figure 4.

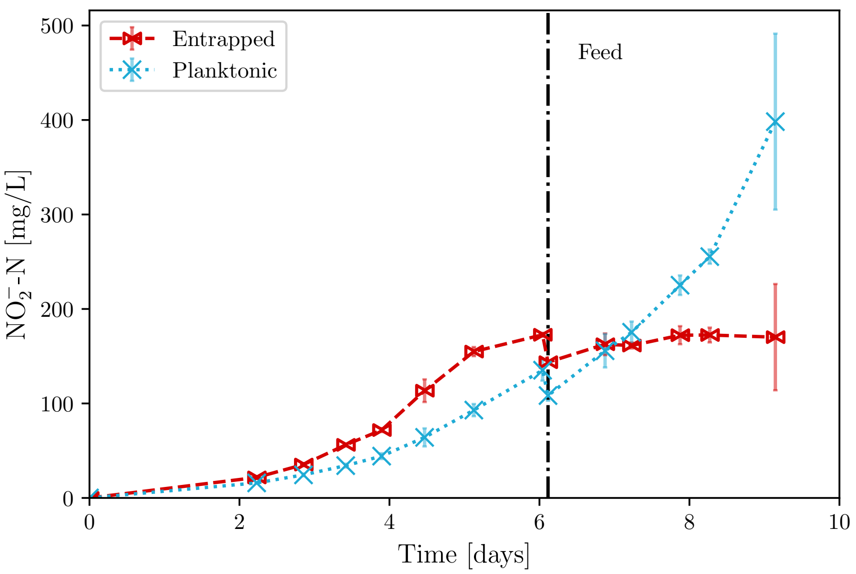


**Fig. S10**: Nitrite concentrations for entrapped and suspended AOB. The timing of the feed led to an increased lag phase in the entrapped culture.

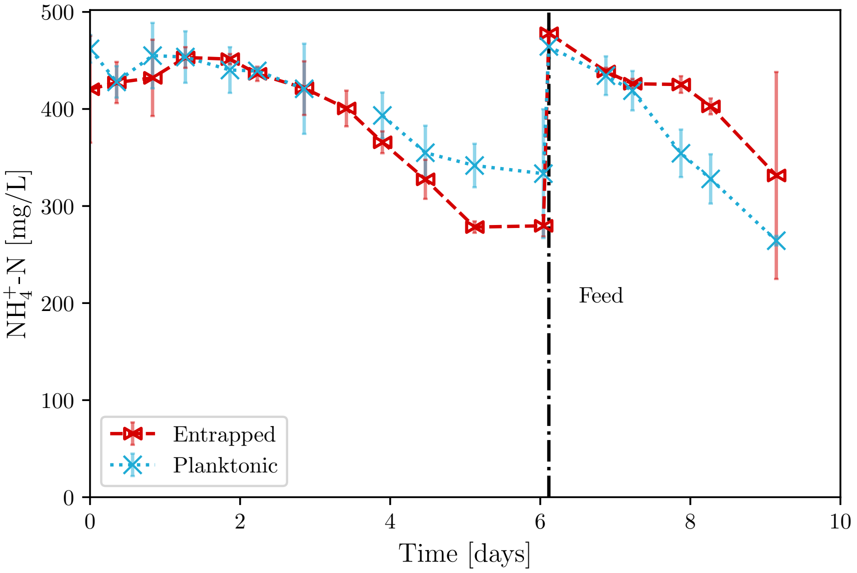


**Fig. S11**: Ammonium concentrations for entrapped and suspended AOB corresponding to Figure S10. Note that there is a higher variability and baseline offset in the cation signal due to HEPES interference (see Section 2.5).

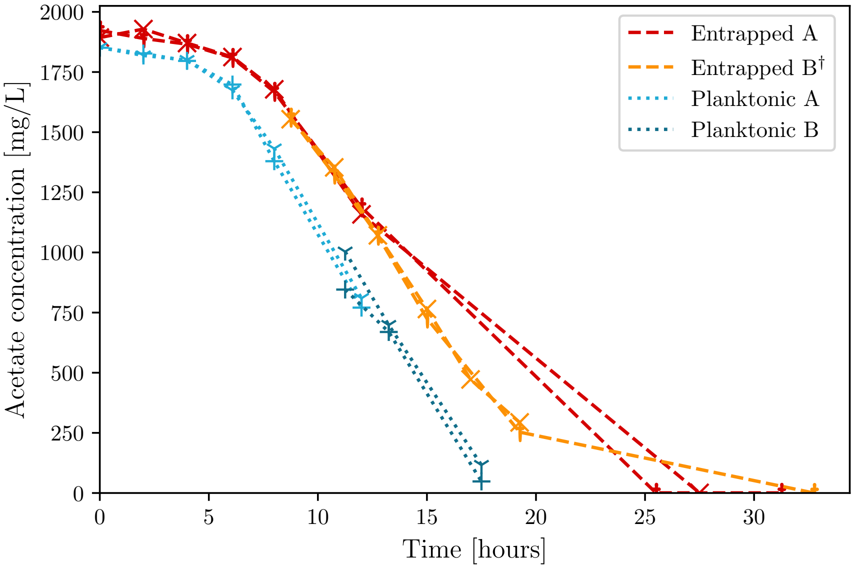


**Fig. S12**: Acetate concentrations for entrapped and planktonic DEN. Groups A and B are duplicates shifted by 12 h to obtain a full metabolite profile. Data of entrapped group B† were time-shifted to compensate for a lag phase induced by storage in the fridge for the first 12 h. This behaviour was not observed in the planktonic replicates of group B.

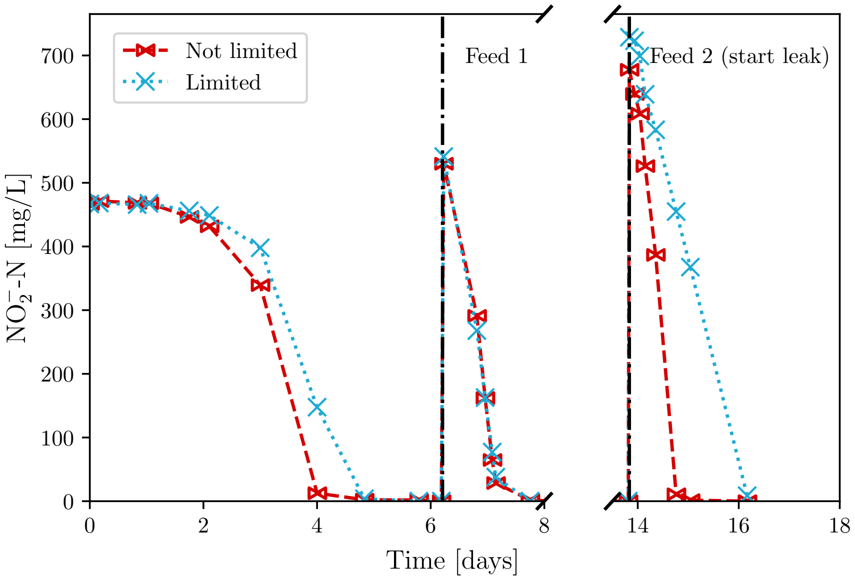


**Fig. S13**: Nitrite concentrations for a reactor limited and not limited by a leak in the air line for suspended NOB. The rate-limited reactor clearly shows a linear (and slower) uptake of nitrite, indicating that no oxygen limitations were present during normal operation.

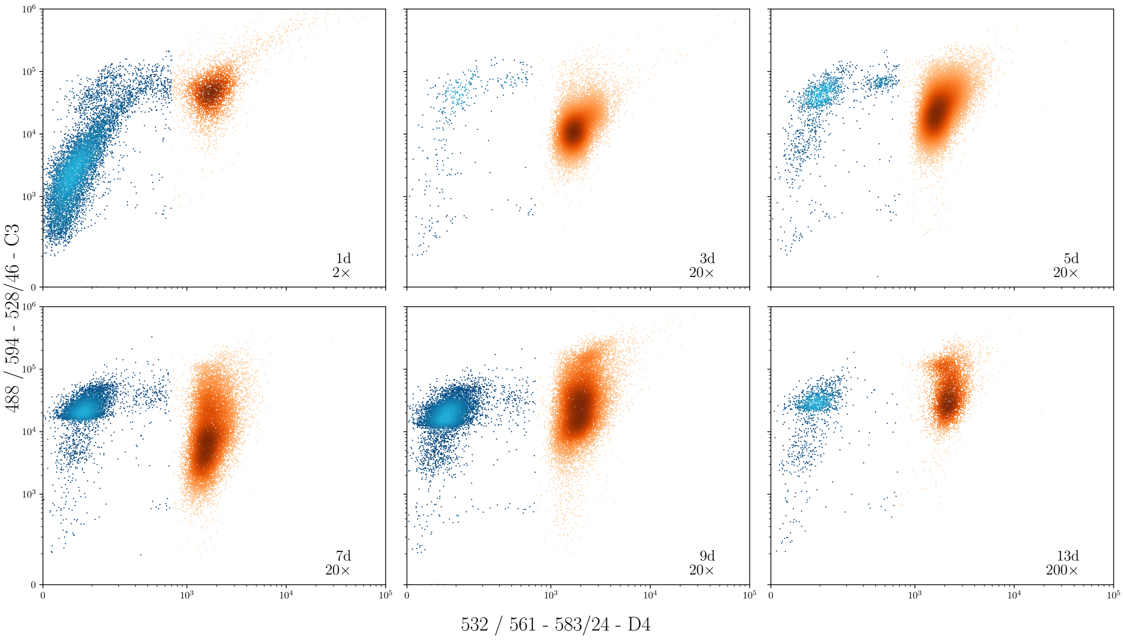


(a)

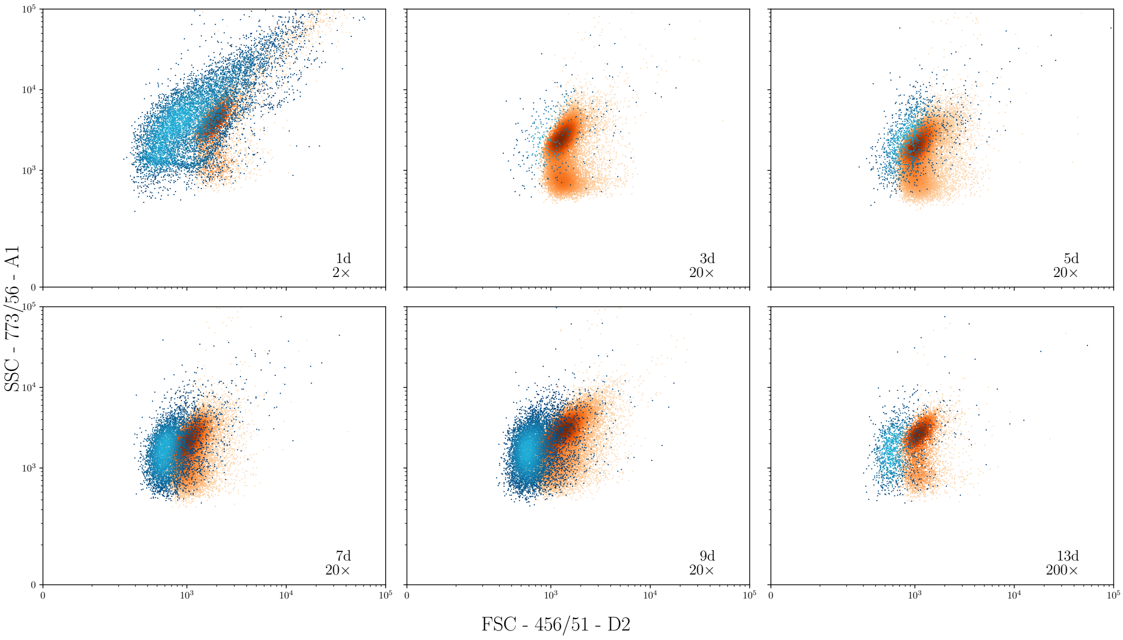


(b)

**Fig. S14**: Flow cytometric assays over time of the SYTO9-stained supernatant of an entrapped coculture reactor run. mScarlet-I positive (DEN) and negative (nitrifiers and DEN not expressing the protein) events are depicted in orange and blue respectively. (a) shows fluorescent channels for SYTO9 (C3) and mScarlet-I (D4), (b) indicates the forward (FSC) and side (SSC) scatter profiles. Clear growth of both populations is visible over time, with a distinct fingerprint in the FSC/SSC channels. Timing and dilution factors are also indicated in the plot.

#### S.2.3 Coculture timelapse

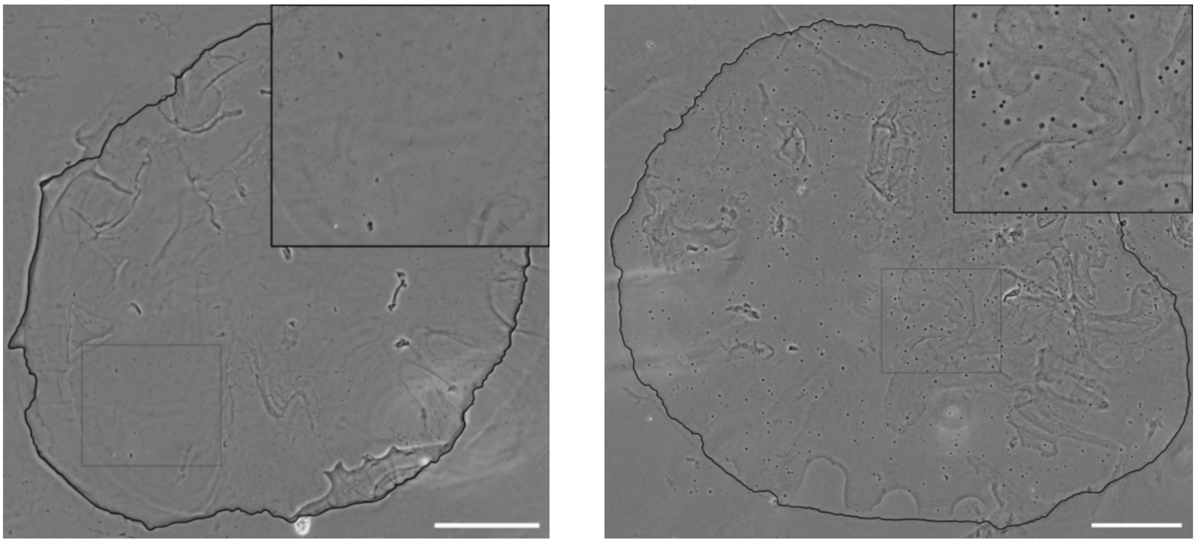

(a) (b)

**Fig. S15**: Phase-contrast images of cryosectioned coculture beads after 0 (a) and 8 (b) days of incubation. Scale bar indicates 200 µm. The top right inset images are a 2X zoom of the area indicated by the gray rectangle and have the same absolute size in both images. The outline of the sections have been enhanced to improve visibility. Colony formation in the right image is homogeneous and clearly present when compared to the initial timepoint on the left.

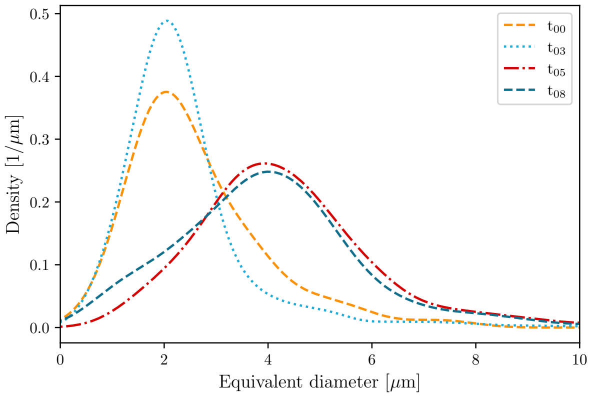


**Fig. S16**: Kernel density estimations of the colony size distribution inside alginate beads over time, as determined by the EUBMIX-FAM signal of FISH-stained cryosectioned beads.

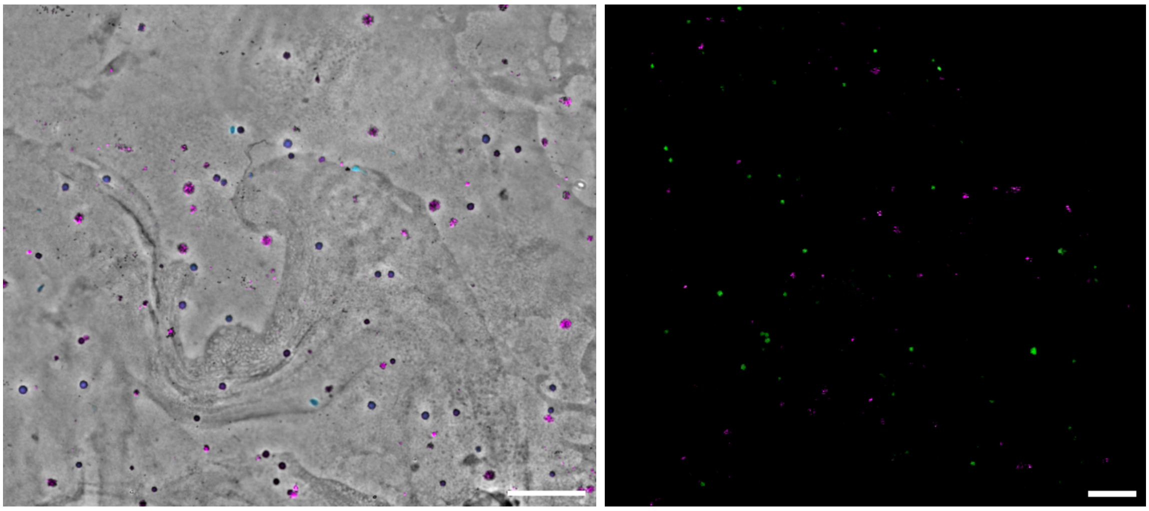

(a) (b)

**Fig. S17**: Cryosectioning-FISH of alginate beads after 8 days of incubation. a) Fluorescence signal layered on top of a phase contrast image. Cyan shows NEU-Atto390 signal targeting AOB, and magenta NIT3-Cy3 targeting NOB (and containing the overlapping mScarlet-I signal from DEN). b) Confocal image with EUBMIX probe (stains all bacteria) indicated in green and magenta coming from NIT3-Cy5 (showing only NOB). Scale bars indicate 50 µm. Colonies are homogeneously distributed, the species themselves are not.

#### S.2.4 Coculture Ion Chromatography (IC) results

***Entrapped experiment***

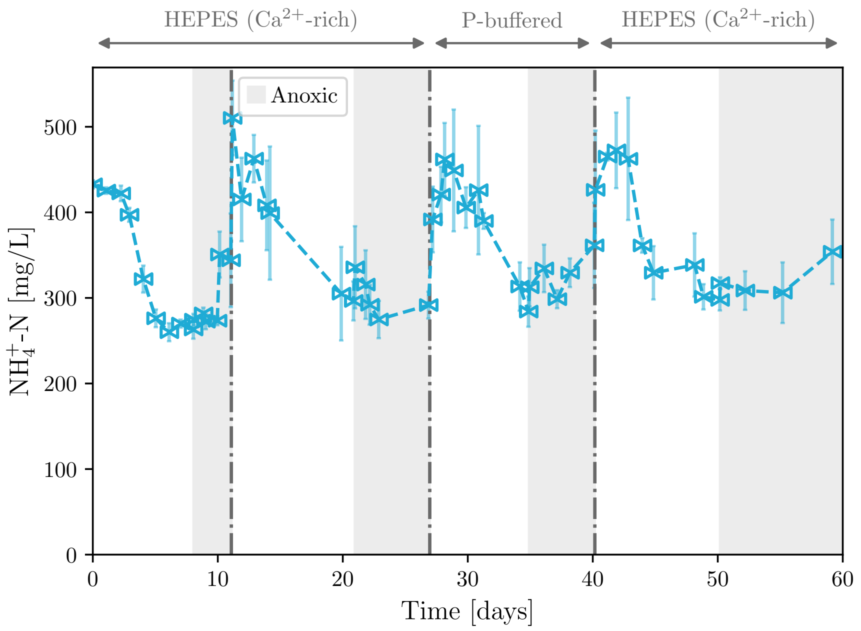


**Fig. S18**: Ammonium concentrations over time of the coculture in entrapped conditions, corresponding to Figure 8. The dash-dotted lines indicate supernatant removal and resuspension in fresh medium. Note that there is a higher variability and a baseline offset in the cation signal due to HEPES interference (see Section 2.5).

***Suspended experiment (planktonic reference)***

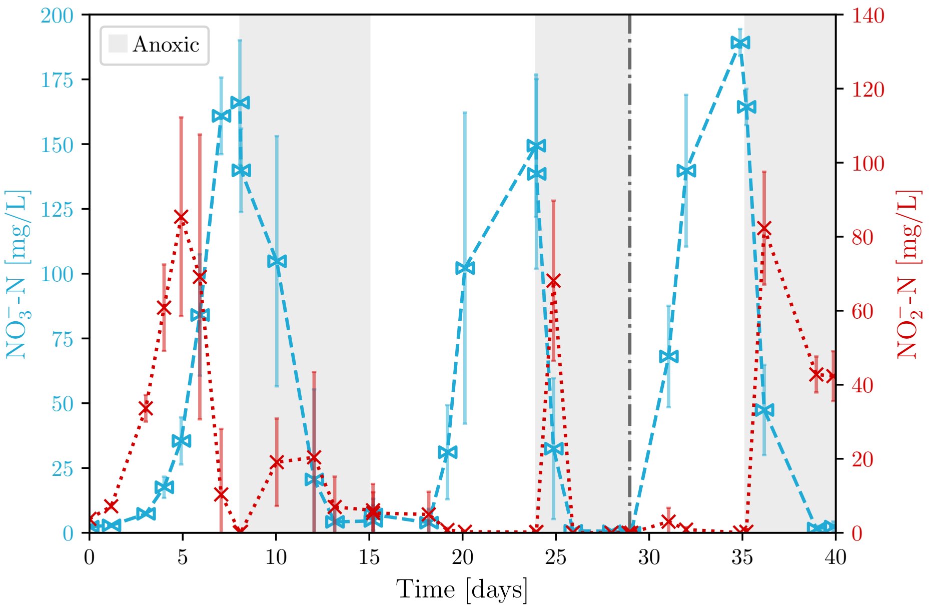


**Fig. S19**: Nitrite (×) and nitrate (⋈) concentrations over time of the planktonic coculture. The dash-dotted line indicates supernatant removal and resuspension in fresh medium.

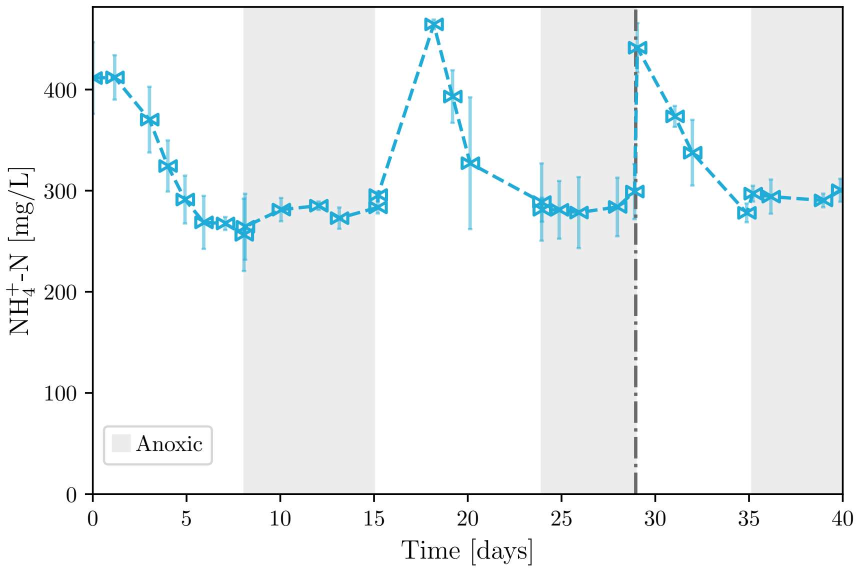


**Fig. S20**: Ammonium concentrations over time of the planktonic coculture, corresponding to Figure S19. The dash-dotted line indicates supernatant removal and resuspension in fresh medium. Note that there is a higher variability and a baseline offset in the cation signal due to HEPES interference (see Section 2.5).

#### S.2.5 Colony release

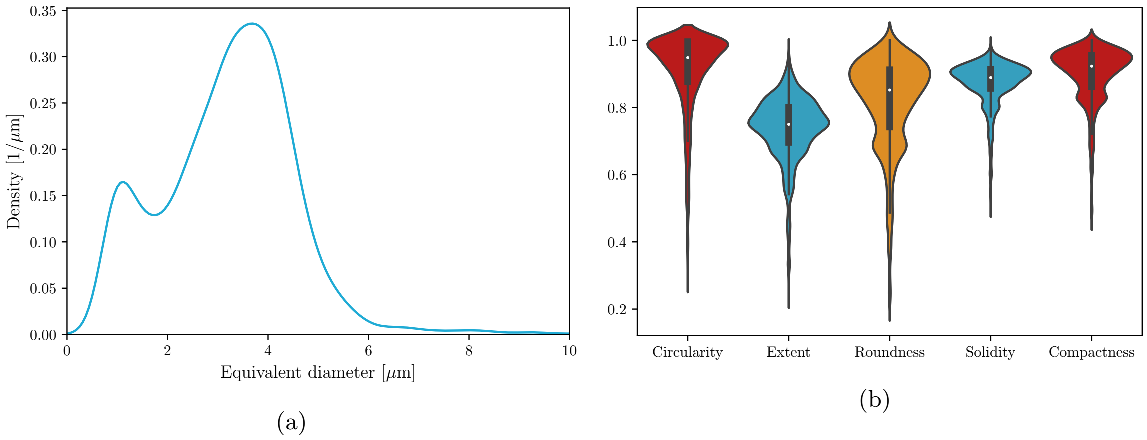


**Fig. S21**: Particle size analysis of the released microcolonies in suspension, based on the SYTO9 signal of pictures taken 9 days after alginate dissolution. a) Shows the kernel density estimation b) Indicates the corresponding particle characteristics.

#### S.2.6 *De novo* flocs over time

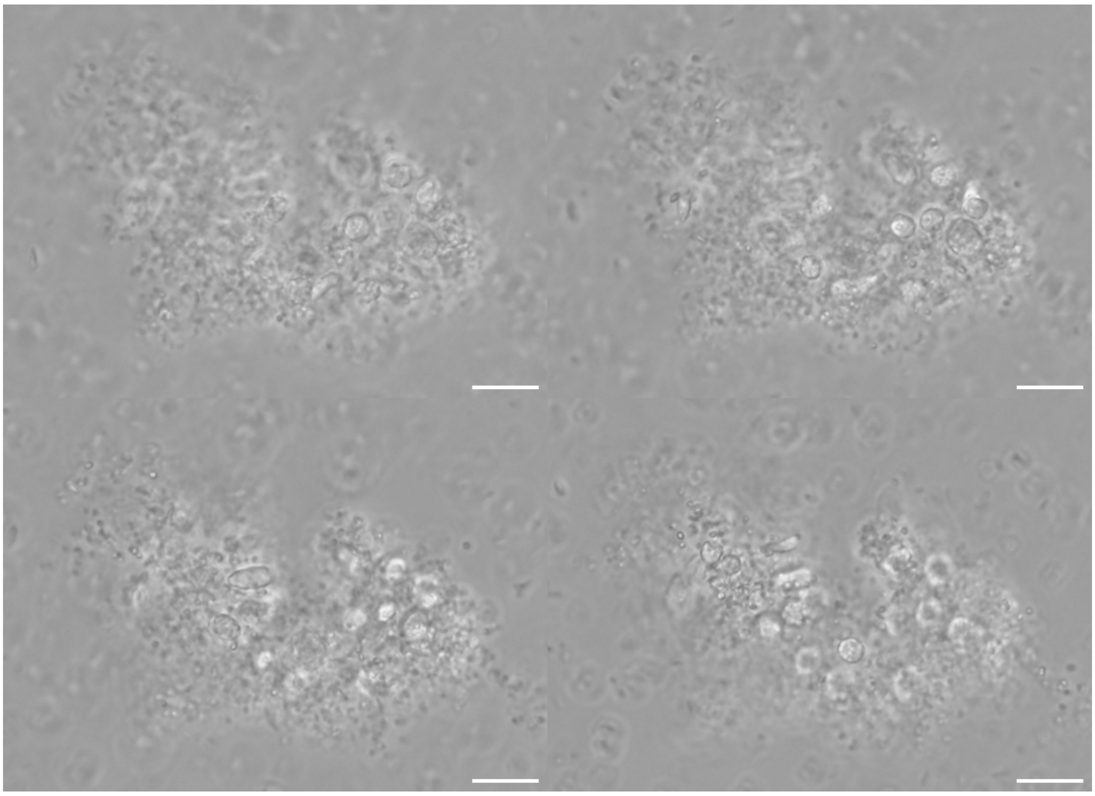

**Fig. S22**: Montage of phase-contrast microscopic images at four different Z positions, showing the presence of the released microcolonies in a DEN matrix 5 days after centrifugation and resuspension in $Ca^{2+}$-rich HEPES-buffered medium. Scale bars indicate 10 µm.

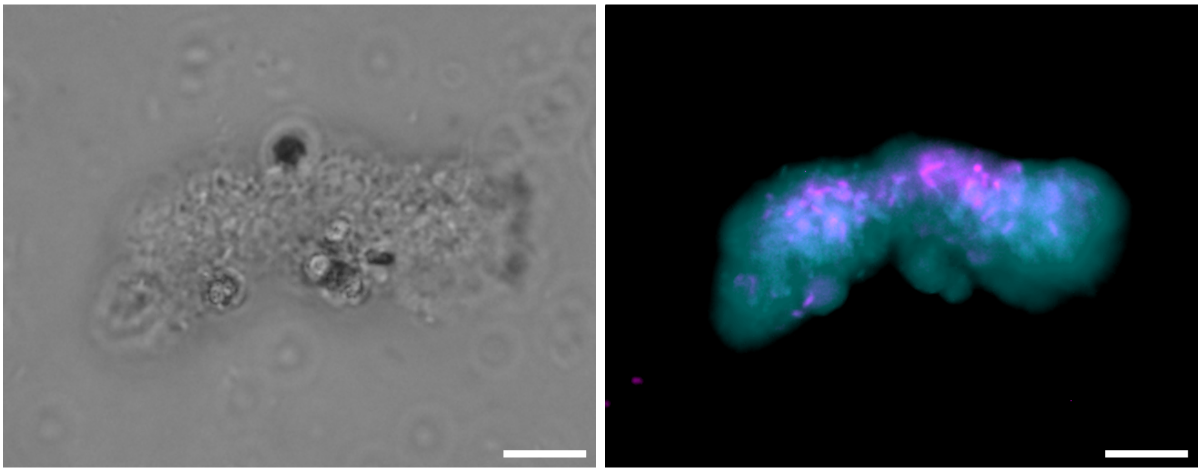

(a) (b)

**Fig. S23**: Suspended clustering example 8 days after centrifugation and resuspension in $Ca^{2+}$-rich HEPES-buffered medium. (a) shows a phase-contrast image, with (b) the corresponding fluorescent signals of SYTO9 in cyan and mScarlet-I in magenta. Scale bars indicate 10 µm.

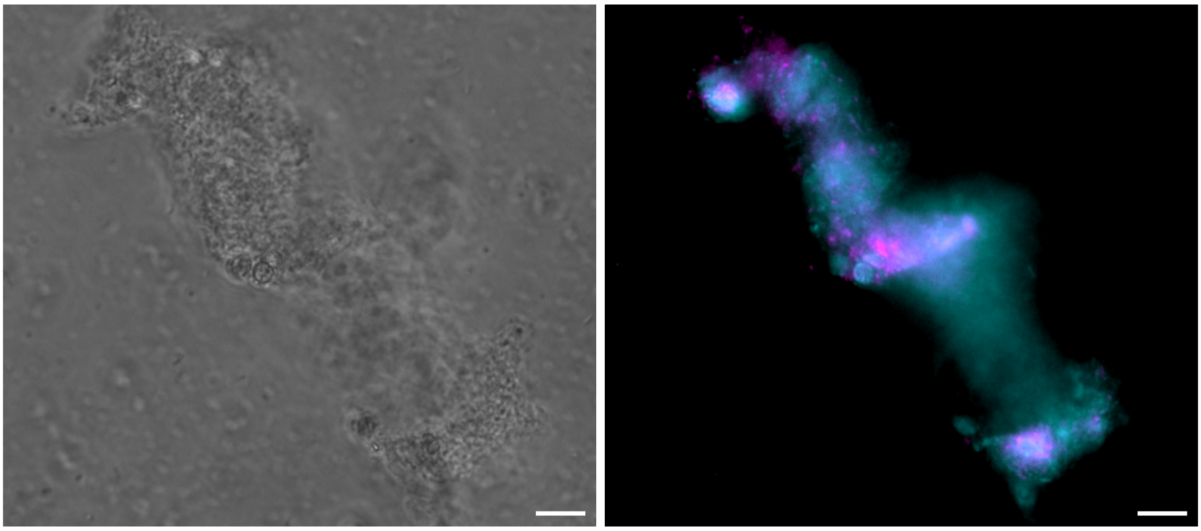

(a) (b)

**Fig. S24**: Suspended clustering example 8 days after centrifugation and resuspension in $Ca^{2+}$-rich HEPES-buffered medium. (a) shows a phase-contrast image, with (b) the corresponding fluorescent signals of SYTO9 in cyan and mScarlet-I in magenta. Scale bars indicate 10 µm.

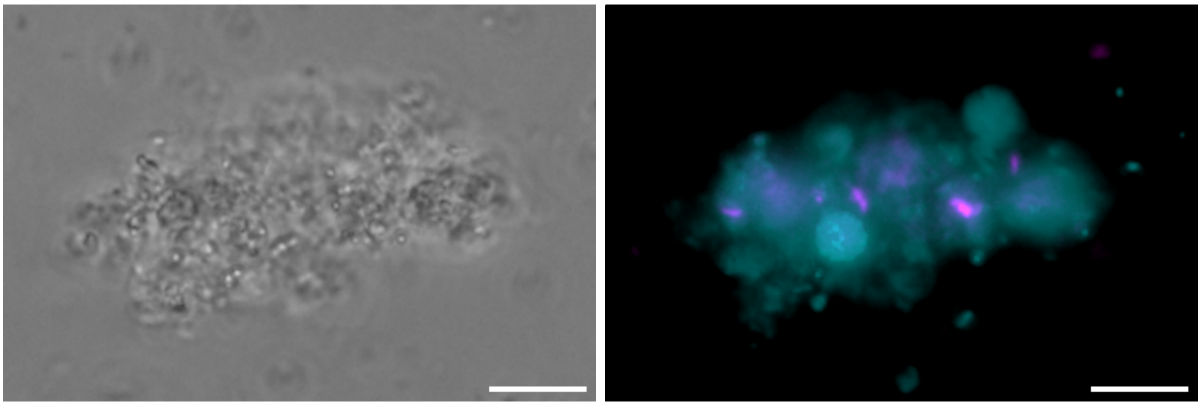

(a) (b)

**Fig. S25**: Less dense suspended clustering example 8 days after centrifugation and resuspension in $Ca^{2+}$-rich HEPES-buffered medium. (a) shows a phase-contrast image, with (b) the corresponding fluorescent signals of SYTO9 in cyan and mScarlet-I in magenta. Scale bars indicate 10 µm.

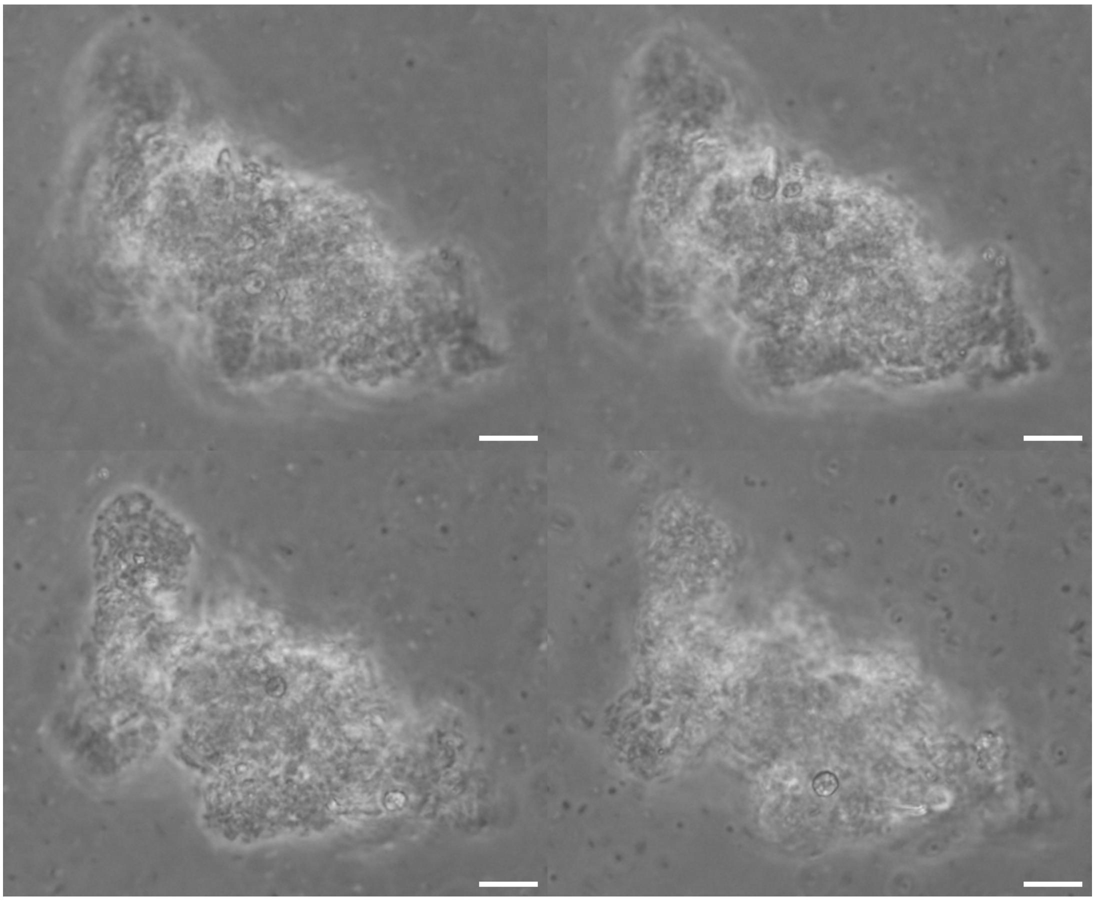

**Fig. S26**: Montage of phase-contrast microscopic images at four different Z positions, showing the presence of the released microcolonies in a DEN matrix after 10 days of incubation. Scale bars indicate 10 µm.

#### S.2.7 Centrifugation effects

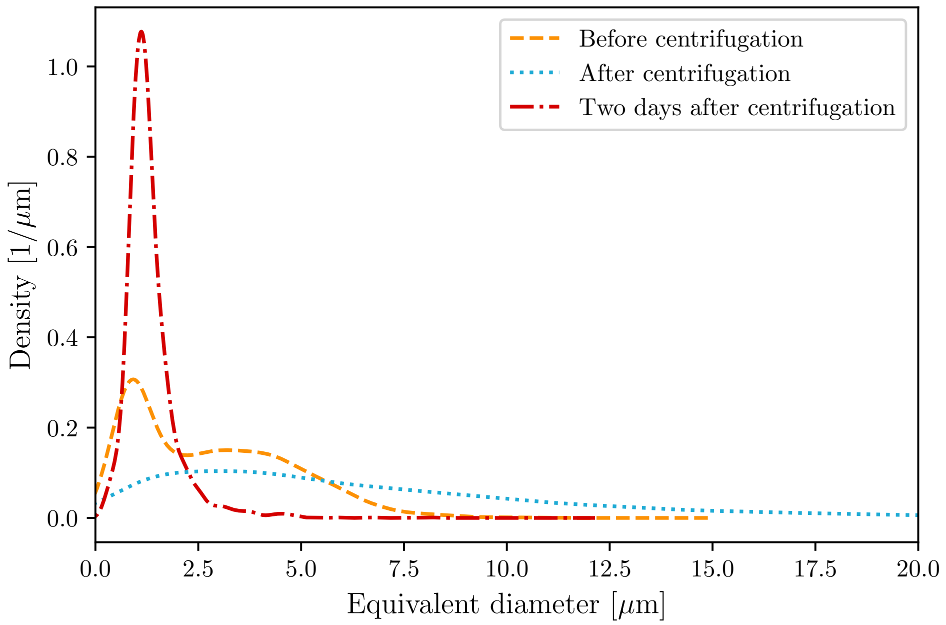


**Fig. S27**: Particle size distributions of *Azoarcus* clusters before and after centrifugation (based on mScarlet-I signal) in the suspended coculture reference experiment. The compaction after centrifugation is reversible. It should be noted that this last sample was taken after a switch to oxic conditions.

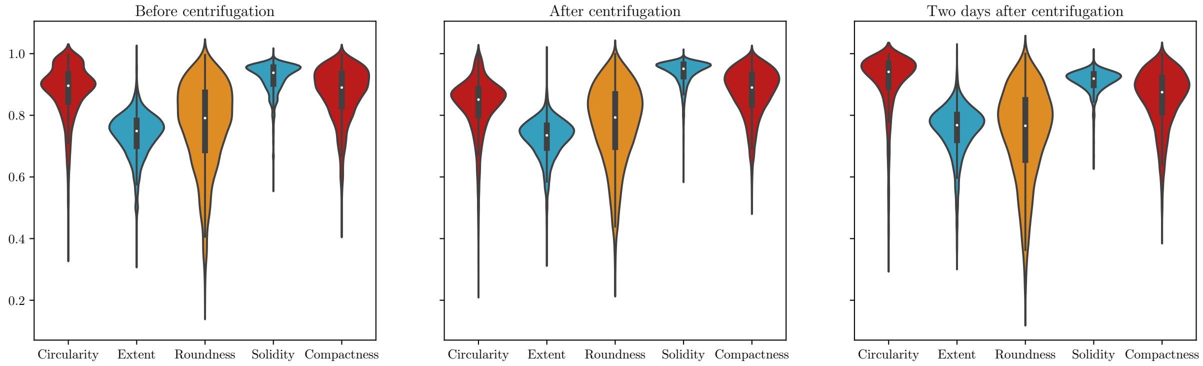


**Fig. S28**: Particle characteristics corresponding to Figure S27.

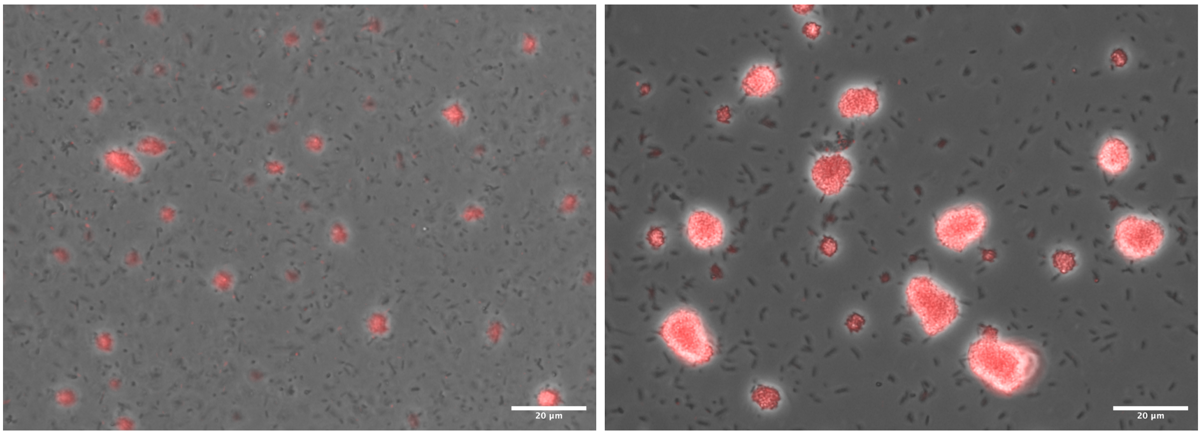


(a) (b)

**Fig. S29**: Example microscopic images of the planktonic coculture reference (a) before and (b) after centrifugation, corresponding to Figure S27. The overlay of the mScarlet-I signal (red) with the phase-contrast channel clearly indicates the centrifugal compaction of the *Azoarcus* clusters. No nitrifier clusters were observed.
